# HPLC Method to Resolve, Identify and Quantify Guanine Nucleotides Bound to the GTPase Ras

**DOI:** 10.1101/2021.06.28.450218

**Authors:** Jonathan P. Hannan, G. Hayden Swisher, Justin G. Martyr, Nicholas J. Cordaro, Annette H. Erbse, Joseph J. Falke

## Abstract

The Ras superfamily of small G proteins play central roles in diverse signaling pathways. Superfamily members act as molecular on-off switches defined by their occupancy with GTP or GDP, respectively. *In vitro* functional studies require loading with a hydrolysis-resistant GTP analogue to increase the on-state lifetime, as well as knowledge of fractional loading with activating and inactivating nucleotides. The present study describes a method combining elements of previous approaches with new, optimized features to analyze the bound nucleotide composition of a G protein loaded with activating (GMPPNP) or inactivating (GDP) nucleotide. After nucleotide loading, the complex is washed to remove unbound nucleotides then bound nucleotides are heat-extracted and subjected to ion-paired, reverse-phase HPLC-UV to resolve, identify and quantify the individual nucleotide components. These data enable back-calculation to the nucleotide composition and fractional activation of the original, washed G protein population prior to heat extraction. The method is highly reproducible. Application to multiple HRas preparations and mutants confirms its ability to fully extract and analyze bound nucleotides, and to resolve the fractional on- and off-state populations. Furthermore, the findings yield a novel hypothesis for the molecular disease mechanism of Ras mutations at the E63 and Y64 positions.

## **1.** INTRODUCTION

The Ras superfamily of small GTP binding and hydrolyzing proteins (GTPases) act as nucleotide dependent regulators in a variety of signal transduction pathways, including those key to cellular activities such as proliferation, migration, differentiation and apoptosis [1–5]. All of these proteins act as binary molecular switches which cycle between a GTP-bound active state and a GDP-bound inactive state [6–9]. In addition to the activating or non-activating nucleotide, the active site cleft also contains a Mg^2+^ ion essential for high affinity nucleotide binding and for GTP hydrolysis to GDP. The Ras superfamily contains greater than 150 members in humans, where the major subfamilies include Ras, Rho, Rab, Rap, Arf, Ran, and Rheb [10].

GTP binding drives a conformational change in Ras superfamily members that primarily alters the structure and/or dynamics of the switch I and II regions, thereby regulating effector docking surfaces to increase the binding affinity of effector proteins (reviewed in [11] . The resulting GTP:Ras:effector complexes trigger a number of essential signaling events in normal cell processes and in disease states, including lipid and protein kinase cascades at cell membrane surfaces. Well-characterized effectors for superfamily members include the phosphoinositide 3-kinases (PI3Ks), the Raf kinases, the Ral guanine nucleotide dissociation stimulator (RalGDS) and the putative tumor suppressor, NORE1 [5, 12, 13].

The four prototypical isoforms of the canonical, founding subfamily are collectively termed "Ras" (HRas, NRas and KRas4A and KRas4B). The biophysical parameters of these isoforms are crucial to their functions and pathologies and illustrate widespread features of Ras superfamily members. The affinities of canonical Ras isoforms for the key guanine nucleotides, (GTP and GDP) is in the picomolar to nanomolar range [14]. A consequence of this high affinity is a relatively slow dissociation rate of bound nucleotide, with a reported half-life of one or more hours in the presence of physiological levels of Mg^2+^ ion [15]. Similarly, nucleotide exchange of GTP for GDP (or vice-versa) is also slow, on a timescale exceeding one hour. The activated GTP:Ras state is terminated by the hydrolysis of GTP to GDP which eliminates the γ-phosphate of GTP. For isolated GTP:Ras, intrinsic GTP hydrolysis is slow, with a reported lifetime of ∼1.4 hrs at room temperature or ∼25 min at 37 °C [16–19]. By contrast, cell signaling pathways typically require rapid on-off switching on a much more rapid timescale. Thus, many signaling pathways provide GTPase activating proteins (GAPs) to speed GTP hydrolysis and conversion to the inactive state, as well as GDP-GTP exchange factors (GEFs) to speed restoration of the GTP-stabilized active state.

The four canonical Ras isoforms are strongly linked to oncogenesis. According to NIH / National Cancer Institute, Ras mutations are present in at least 30% of human cancers (https://www.cancer.gov/research/key-initiatives/ras). The COSMIC database indicates there are three cancer-associated mutational hotspots on the conserved Ras GTPase domain at positions Glycine 12, Glycine 13 and Glutamine 61 [20]. At these hotspot positions side chain substitutions are cancer-linked due to a compromised ability to intrinsically hydrolyze GTP to GDP and/or compromised interactions with GAP proteins [20–24] and reviewed in [25, 26]. Thus, the mutation retains a higher than native fraction of the Ras population in the GTP-occupied active state, thereby yielding superactivation that can drive or support oncogenesis and tumorigenesis. Less well understood are cancer-linked mutations at numerous positions outside the three hotspots. At these non-hotspot positions, one or a few specific side chain substitutions are cancer-linked ((https://cancer.sanger.ac.uk). In general, the molecular cancer mechanisms of these non-hotspot mutations have yet to be discovered.

As a result of their central role in native signaling pathways and in pathologies, Ras proteins have been extensively investigated from both a structural and biochemical perspective. The best characterized is HRas, with numerous three-dimensional structures of both wild-type and disease-associated variants having been determined by X-ray crystallography and nuclear magnetic resonance (NMR) [11, 27]. These include structures of HRas in both inactive and active states, loaded with GDP or a hydrolysis-resistant GTP analog such as guanosine 5′-[β,γ-imido]triphosphate (GMPPNP), respectively (**Figure 1**). Notably, the structure of GMPPNP is identical to that of GTP except that the oxygen atom bridging the β-phosphate to the γ-phosphate is replaced by a secondary amine group (**Figure 2**). As a result, GMPPNP is an excellent surrogate for GTP and activates Ras proteins in a manner similar to GTP [28]. Also available are co-complex structures of HRas bound to effector proteins, including HRas:PI3Kγ, HRas:Raf Ras Binding Domain, HRas:grb14 Ras Associating Domain, and HRas:NORE1A [29–32].

**Figure 1.**
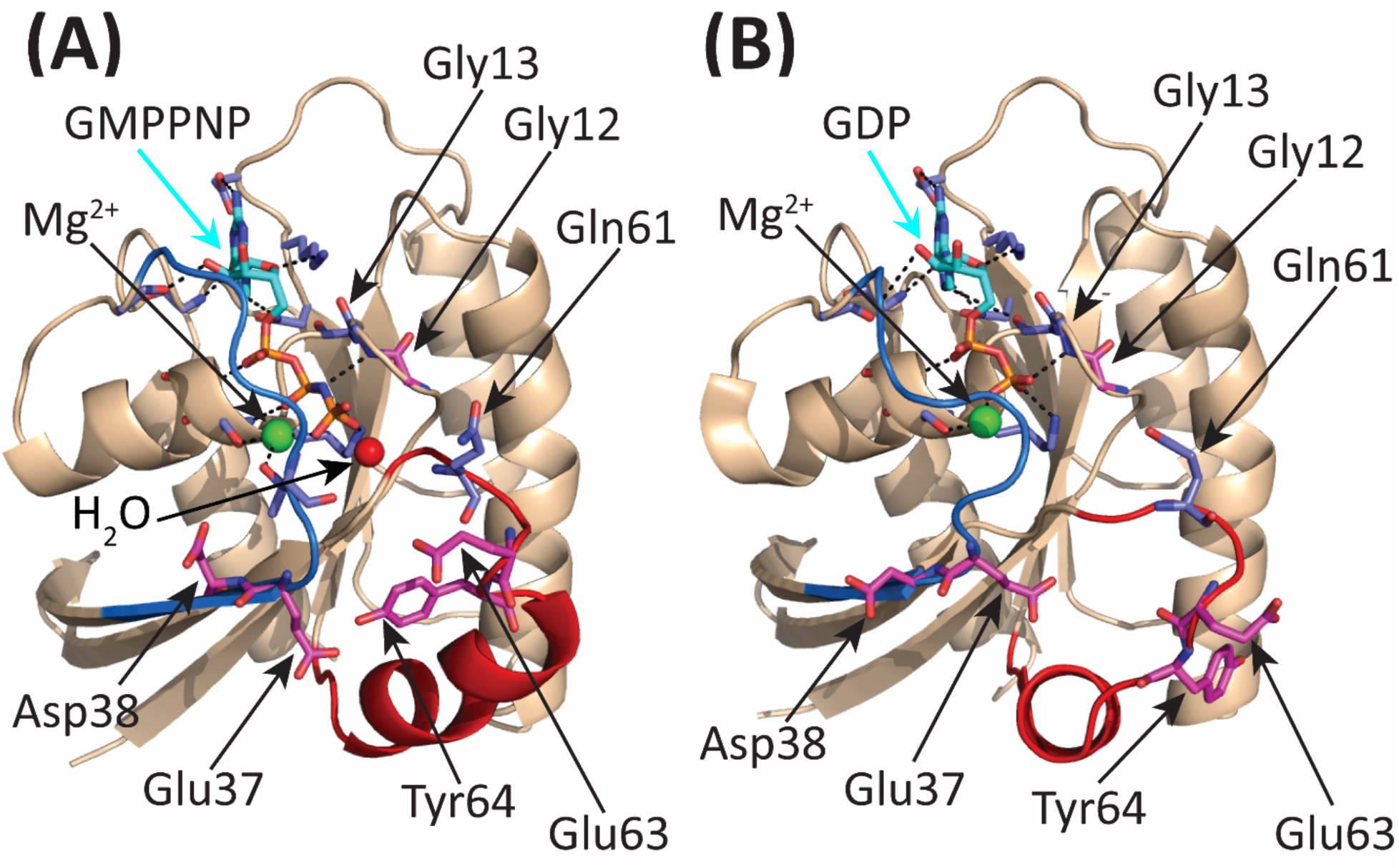
Structures of active and inactive HRas loaded with GMPPNP and GDP, respectively. **(A)** Cartoon representation of active HRas loaded with the hydrolysis-resistant GTP analog, GMPPNP (PDB ID: 5P21) [6]. Highlighted are Switch I (blue backbone and Switch II (red backbone), the bound GMPPNP nucleotide (colored atoms with cyan carbons), the active site Mg^2+^ ion (green sphere), the water molecule closest to the site of nucleophilic attack during GTP hydrolysis (red sphere), native side chains mutated in the present study (labeled), and native side chains involved in nucleotide coordination (unlabeled with dashed coordination). **(B)** Cartoon representation of inactive HRas loaded with GDP (PDB ID: 4Q21) [7]. Color coding same as **(A)**.

**Figure 2.**
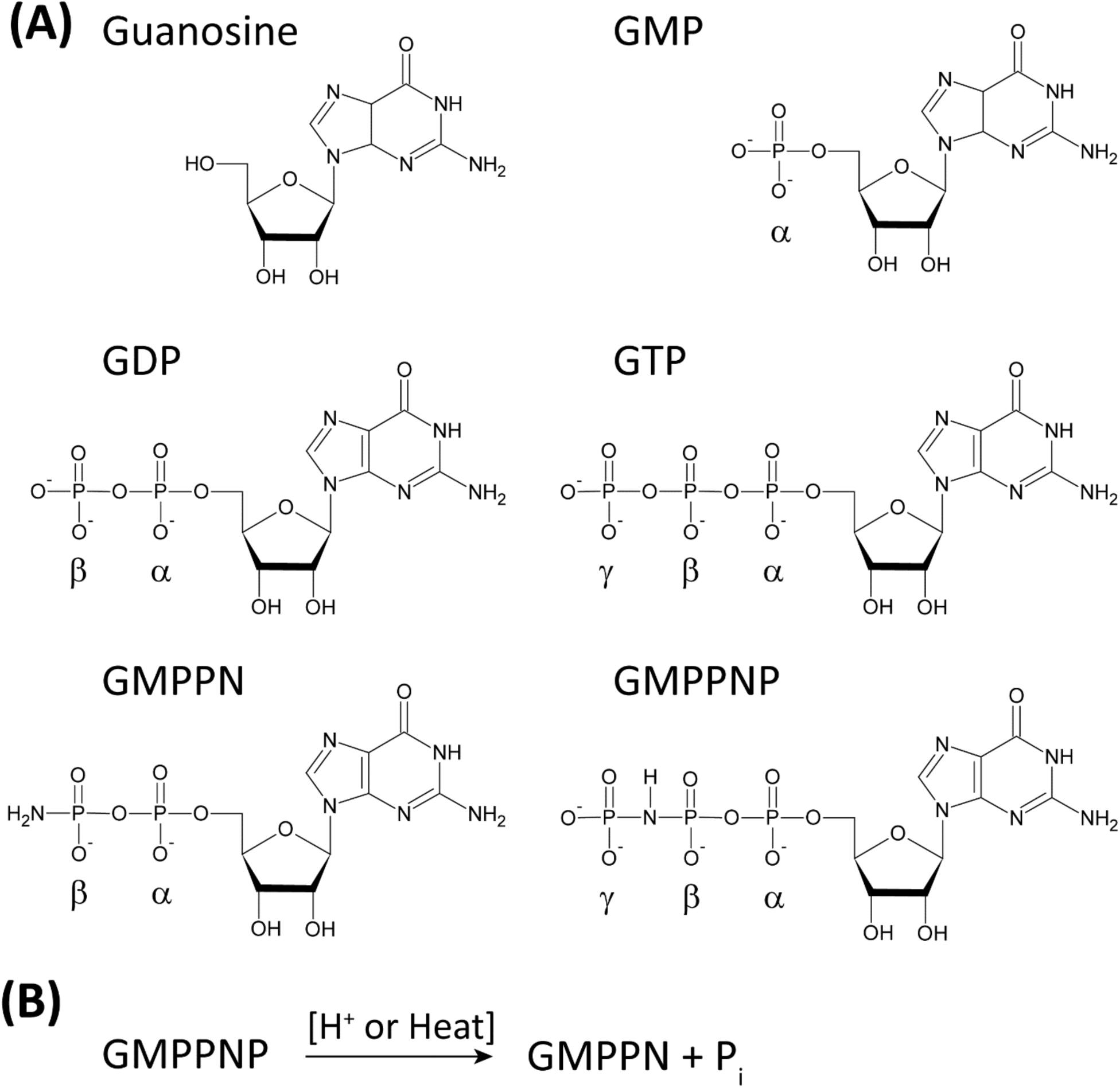
Guanosine and the guanine nucleotides utilized herein. **(A)** Structural formulae of guanosine and the non-cyclic guanine nucleotides monitored in the present study: guanosine-5′-monophosphate (GMP); guanosine-5′-diphosphate (GDP); guanosine-5′-triphosphate (GTP); guanosine-5′-(β-amino)-diphosphate (GMPPN); and guanosine 5′-[β,γ-imido]triphosphate (GMPPNP) **(B)** The GTP analog GMPPNP is strongly resistant to hydrolysis by Ras but can be hydrolyzed to GMPPN by acidic conditions or high temperatures.

For quantitative *in vitro* studies of Ras function, it is crucial to measure the fractions of the Ras population loaded with activating and inactivating nucleotides. The goal of the present study is to develop a rigorous, quantitative HPLC procedure that can resolve and quantify multiple nucleotides bound to Ras. A given sample of Ras will often be loaded with a mixture of different guanine nucleotides. For example, recombinant Ras isolated from cells is a mixture of GTP:Ras and GDP:Ras, while purified Ras loaded *in vitro* with GMPPNP may be a mixture of GMPPNP:Ras with residual GTP:Ras and GDP:Ras from the cell. Here we describe a novel HPLC method that provides resolution, identification and quantification of the guanine nucleotide subpopulations bound to the GTPase. The approach employs GMPPNP rather than GTP to stabilize the active state, both because GMPPNP is resistant to intrinsic Ras hydrolysis, and because GMPPNP is quantitatively converted by heating to the product guanosine-5′-(β-amino)-diphosphate (GMPPN or GppNH_2_) with the accompanying release of inorganic phosphate (**Figure 2**) [28, 33]. The fortuitous hydrolysis of GMPPNP to an easily identified product, together with the heat resistance of GTP and GDP, allow use of a heat denaturation step to quantitatively extract the tightly bound nucleotide population from a washed Ras preparation while precipitating the protein for removal [34]. The resulting isolated nucleotide population is then analyzed by reverse-phase HPLC-UV to quantify its individual guanine nucleotide components. Ultimately, the analysis yields the nucleotide loading of the original HRas population (prior to heating) as well as its fractional distribution between the activated (loaded with GMPPNP or GTP) and inactivated (loaded with GDP) signaling states. Notably, application of the method to multiple HRas preparations and mutants illustrates its ability to reproducibly measure their nucleotide loading parameters, and thus their signaling state distributions. Validation of the method is provided by excellent agreement between the total nucleotide concentration measured by HPLC-UV and the total G protein concentration measured independently both by the G protein UV deconvolution method and by quantitative amino acid analysis [34], as expected for the well-established 1:1 nucleotide:protein stoichiometry of the complex. Finally, the findings unexpectedly reveal a novel hypothesized molecular disease mechanism for mutations at the E63 and Y64 positions.

## 2. MATERIALS AND METHODS

### 2.1. Recombinant HRas expression and purification

The plasmid used to express standard and mutant HRas proteins in *E. coli* is a modification of a previously described HRas expression plasmid [35–37]. The modified plasmid expresses HRas amino acids 1-184 with two site-directed Cys substitutions (C118S/C181S) and one truncation (Δ185-189) to eliminate all surface Cys residues except one (C184), thereby minimizing artifacts due to promiscuous disulfide bond formation, and facilitating homogeneous membrane anchoring or spectroscopic labeling when desired [35]. The resulting modified HRas, termed "standard" HRas, acted as our ‘wild-type’ surrogate, designated as HRas hereafter, and also served as the fixed background for the creation of all additional HRas mutants employed in this study (G12S, E37G, D38E, E63K and Y64G). All mutations were generated by site directed mutagenesis (Agilent Technologies; mutagenesis primers from Integrated DNA Technologies) followed by sequencing of the full HRas gene (GENEWIZ).

Individual HRas proteins were expressed in *Escherichia coli* BL21 DE3 cells based on established protocols [35–38]. Briefly, the desired HRas plasmid was expressed in *E. coli* BL21 DE3 cells at 37 °C followed by induction with 0.5 mM isopropyl β-d-1-thiogalactopyranoside (IPTG) at 18°C for 20 hrs. Cells were then harvested, lysed by sonication, centrifuged to clarify, and HRas was purified via its 6His affinity tag using immobilized metal affinity chromatography (TALON® Metal affinity resin (Takara)) followed by elution using an imidazole step gradient. The resulting, purified HRas was loaded with a mixture of guanine nucleotides (GXP), in this case originating from its *E. coli* expression. Here the GXP nucleotide population was mainly GDP, and a smaller fraction of GTP (since most bound GTP from the cell is hydrolyzed by intrinsic HRas GTPase activity during purification).

### 2.2 Loading HRas with a desired nucleotide

Stable samples of activated and inactivated HRas were generated by exchanging the intrinsic bound GDP/GTP nucleotide population for a non-hydrolyzable GTP analog (GMPPNP) or for GDP, respectively. The exchange was facilitated using EDTA to chelate the Mg^2+^ ion cofactor of the HRas nucleotide binding pocket, thereby destabilizing the bound nucleotide and accelerating the intrinsic exchange reaction by orders of magnitude. Following exchange with ≥ 10-fold molar excess of the desired nucleotide, the exchange reaction was quenched by addition of excess Mg^2+^ to restore the high kinetic and thermodynamic stability of the bound nucleotide population.

During the development of the HRas loading protocol, two different procedures (Loading Protocols I and II, respectively) were compared. Loading Protocol I utilized a methodology adapted from that developed by Martyr, Swisher and Falke [38]. Briefly, affinity purified HRas samples were concentrated and Mg^2+^ was removed via buffer exchange into 40mM HEPES, pH 7.4, 50 mM EDTA, 150 mM NaCl, 2.5 mM Glutathione, 10% glycerol using Vivaspin 500 10kDa MWCO spin concentrators (Sartorious). The remaining Mg^2+^-EDTA chelates and free nucleotides were then removed by dialysis using slide-a-lyzer dialysis cassettes (10,000 MWCO; Thermo-Fisher) while concomitantly facilitating exchange of protein samples into 25 mM HEPES, pH 7.4, 140 mM KCl, 15 mM NaCl, 2.5 mM Glutathione, 10% glycerol. After dialysis, HRas samples were then clarified by centrifugation at 89,000 X *g* for 1 hour at 4**°**C in a Beckman TL-100 ultracentrifuge equipped with a Beckman Coulter TLA 120.2 rotor fixed angle rotor and the concentration of recombinant HRas was then determined by UV deconvolution [34]. Proteins were then incubated with a 10-fold molar excess of the chosen guanine nucleotide (GMPPNP or GDP (AbCam)) for 3 hours at room temperature (22 **°**C) before adding 1 mM excess MgCl_2_ and snap freezing in 50 µl aliquots at a final protein concentration ≤ 220 µM. All samples were stored at -80 **°**C until required for HPLC or biophysical analyses.

For Loading Protocol II, a procedure adapted from that utilized by Smith and coworkers was employed [39]. Affinity purified HRas was buffer exchanged into 25 mM HEPES, pH 7.4, 140 mM KCl, 15 mM NaCl, 2.5 mM Glutathione, 1 mM MgCl_2_ using EconoPac 10DG desalting columns (Bio-Rad). The protein concentration was then determined by a UV deconvolution procedure [34]. Potential residual cellular phosphatase activity was blocked by incubating HRas samples for 10 min at 37 °C with PhosSTOP (Roche). HRas samples were subsequently incubated for 20 min at 37 °C with 10 mM EDTA and a 15X molar excess of either GMPPNP or GDP. After incubation, sufficient MgCl_2_ was added to obtain a final concentration of 2 mM free MgCl_2_. HRas samples were then exchanged into 25 mM HEPES, pH 7.4, 140 mM KCl, 15 mM NaCl, 2.5 mM Glutathione, 10% glycerol using EconoPac 10DG desalting columns (Bio-rad) and concentrated to ∼220 µM using Vivaspin 500 10kDa MWCO spin concentrators (Sartorious). The final protein concentration was verified by UV deconvolution and GXP:HRas samples were then snap-frozen in 55 µl aliquots and stored at -80 **°**C until required[34].

### 2.3 Preparation of washed GXP:HRas complexes

Following nucleotide loading GXP:HRas complexes were washed to remove unbound components and concomitantly buffer exchange the samples into an HPLC sample buffer comprising 25 mM HEPES, pH 7.4, 140 mM KCl, 15 mM NaCl. Two buffer exchange procedures were developed to wash the nucleotide-HRas complexes for different applications (designated Wash Protocol A and Wash Protocol B, respectively). Wash Protocol A was employed when extraction and HPLC quantification of the bound nucleotides was desired and protein quantification was not needed, while Wash Protocol B was employed when extraction and quantification of bound nucleotides was required with parallel quantification of HRas protein by UV deconvolution [34]. Both methods utilized repetitive cycles of dilution of GXP-HRas with HPLC sample buffer, followed by ultrafiltration back to the predilution volume, yielding a net 50,000-fold effective dilution of any residual components not tightly bound to HRas.

For Wash Protocol A, 100 µl of GXP:HRas sample (obtained by thawing two aliquots of stored protein on ice) was buffer exchanged into HPLC sample buffer using Vivaspin 500 spin concentrators, 10 KDa MW cutoff membrane (Sartorious) in conjunction with a Beckman Coulter microfuge 18 centrifuge (11,000 X *g*; 4 °C, room temp buffer). A total of five successive exchange steps were carried out facilitating the removal of excess free GXP nucleotide and undesired residual components (glutathione, glycerol) of the protein prep, yielding a net 50,000-fold effective dilution of these unbound components. After buffer exchange, the GXP:HRas sample was diluted to 20-50 µM concentration in HPLC sample buffer, before being processed for HPLC analysis as detailed below in **Section 2.4**.

For Wash Protocol B an adaption of a method previously described by Swisher et al., was utilized [34]. First 100 µl of GXP:HRas was thawed on ice and then buffer exchanged into HPLC sample buffer using Vivaspin 500 spin concentrators, 10 KDa MW cutoff membrane (Sartorious) as described above for Wash Protocol A. After buffer exchange, GXP:HRas samples were diluted in HPLC sample buffer to give a final volume of 340-400 µl (20-50 µM protein), and then subjected to ultracentrifugation at 89,000 X *g* for 40 min at 22 **°**C in a Beckman TL-100 ultracentrifuge equipped with a Beckman Coulter TLA 120.2 rotor fixed angle rotor to remove any protein aggregates. The HRas protein concentrations of the washed GXP:HRas samples were then determined by UV deconvolution as previously outlined [34].

### 2.4 Extraction of nucleotides from nucleotide-HRas complexes for HPLC analysis

To extract the nucleotides from a washed nucleotide-HRas complex, a heat-extraction procedure was used to denature the protein, release the bound nucleotide, and the precipitated protein removed by centrifugation. HRas samples were heated at 95 °C for 6 min using a MiniAmp thermocycler (Applied Biosystems), resulting in protein denaturation, release of associated GXP nucleotides, as well as quantifiable levels of nucleotide hydrolysis (**Section 2.7**). After heating, denatured/precipitated protein was pelleted by centrifuging samples at 11,000 X *g* for 10 min at room temperature using a Beckman Coulter microfuge 18 centrifuge, and the supernatant containing the extracted nucleotide mixture was removed and further clarified by passing through a pre-rinsed 0.20 µm Advantage PVDF MicroSpin centrifuge filter at 11,000 X *g* at room temperature for 10 min using a Beckman Coulter microfuge 18 centrifuge.

To verify that nucleotides did not bind to the pre-rinsed PVDF filters, control experiments quantified stock solutions of GMPPNP/GMPPN, and separately, GDP at concentrations ranging of 60 µM, 30 µM, 15 µM, and 7.5 µM by UV spectrometry, both before and after passing through PVDF filtration devices. No changes in nucleotide concentration were observed after passing through the PVDF centrifuge filter (**Supplemental Figure 1**).

### 2.5 Preparation of standard nucleotide samples for HPLC analysis

To serve as references and standards in HPLC analysis of HRas-associated nucleotides, stock solutions of the relevant nucleotides at concentrations of 1 to 20 mM in HPLC sample buffer were prepared from commercial reagents: GTP (Abcam; (>90 % purity)), GMPPNP (Abcam; >95 %)), GDP (Abcam; (>98 %)), GMPPN (hydrolysis product of GMPPNP, JenaBioscience, (≥ 95 %)) and GMP (Abcam, (>98 %)). Stocks were snap-frozen and stored at -80 °C until day of use. Subsequently, the stock was thawed on ice, diluted to a concentration of ∼250 µM in HPLC sample buffer, and the resulting concentration was quantified by its UV absorbance at 252 nm (ε=13.7×10^3^ M^-1^ cm^-1^) [40]. Where indicated, these quantified stocks were used to generate nucleotide mixtures of interest in HPLC sample buffer. One such mixture, used to test the resolution of the HPLC system, contained 40 µM each of GTP, GDP, GMPPNP, GMPPN and GMP. In some cases, nucleotides were also heated to 95 °C for 6 min to model the effect of HRas heat extraction (**Section 2.7**). All nucleotide samples were clarified by passing through a pre-rinsed PVDF centrifuge filter (**Section 2.4**).

### 2.6 HPLC procedure for resolving and quantifying nucleotides

HPLC analysis of standard and HRas-extracted nucleotides was carried out on an Agilent Technology 1260/1290 Infinity HPLC system fitted with an autosampler. The column system employed consisted of a Phenomenex Standard Guard Cartridge System pre-filter fitted to a Phenomenex Gemini 5 µm C18 reverse-phase analytical column (150 X 4.6 mm). The mobile phase consisted of 92.5 mM KH_2_PO_4_, 9.25 mM tetrabutylammonium bromide, pH 6.4, and 7.5% acetonitrile. Each 45 µl sample was injected into the system via a 1000 µl sample loop which had been pre-washed with HPLC sample buffer (25 mM HEPES, pH 7.4, 140 mM KCl, 15 mM NaCl). Samples were kept at room temperature in the autosampler for no longer than 4 hours before injection. All samples were run in the mobile phase at a flow rate of 1.3 ml/min for 9 min at 22 °C. No degradation of individual nucleotides or changes in the ratios of nucleotide mixtures could be detected over the lifetime of the experiments.

To quantify each nucleotide, its UV absorbance was monitored at 252 nm as it emerged from the HPLC column, and then its concentration was determined by comparison with a standard curve generated from a serial dilution of an individual nucleotide (or mixture of nucleotides). Total peak areas (mAU*sec) were quantified for each experimental or standard nucleotide peak.

Standard curves plotted the peak areas of standard nucleotides against their known concentrations, and the best fit straight line (with y-intercept set to 0) determined by linear best fit analysis (Microsoft® Excel® for Microsoft 365)). The slope of the line, which allows correlation between the nucleotide concentration and peak area, was subsequently used to interpolate or extrapolate the concentration of each nucleotide component.

### 2.7 Quantifying the effects of heat treatment on nucleotides

Additional HPLC analyses were carried out to quantify the effects of the heat treatment used to extract nucleotides from HRas. Stock solutions of individual nucleotides (∼250 µM, **Section 2.5**) were prepared and divided into three identical aliquots. One aliquot was placed at room temperature, the second aliquot was heated at 95 °C in a thermocycler for 3 min and then cooled on ice for 2 min, and then allowed to equilibrate to room temperature, and the third aliquot was heated at 95 °C in a thermocycler for 6 min, cooled on ice for 2 min, and then allowed to equilibrate to room temperature. All three samples were then diluted to 60 µM and passed through the PVDF filter (**Section 2.4**) prior to HPLC analysis and quantitation (**Section 2.6**).

### 2.8 Statistics

Data reported for the heat treatment of commercially available guanine nucleotide standards were obtained by averaging the triplicate HPLC-UV measurements of two samples generated independently for each condition. The resulting percentage peak areas for each guanine nucleotide species are reported as the mean ± standard deviation.

For mixtures of nucleotides obtained by heat-extraction from HRas prepared using Wash Protocol A, data were generated by averaging the triplicate HPLC-UV measurements of two samples generated independently for each condition. For mixtures of nucleotides obtained by heat-extraction from HRas prepared using Wash Protocol B, data were generated by averaging the triplicate HPLC-UV measurements of three samples generated independently for each condition. For the latter measurements employing Wash Protocol B, a minimum of three UV spectra were recorded for each independent sample for use in the UV deconvolution procedure to measure the protein concentration [34]. Resulting percentage peak areas for guanine nucleotides and protein concentrations are reported as the mean ± standard deviation.

## 3. RESULTS

### 3.1 HPLC-UV of Guanine Nucleotide Standards

Building on previously reported protocols, we have developed an ion-pair, reverse-phase HPLC procedure to resolve, identify and quantify the major guanine nucleotides bound to HRas [41–53]. The procedure employs a C18 analytical column running a mixed-solvent mobile phase (92.5 mM KH_2_PO_4_, 9.25 mM tetrabutylammonium bromide, pH 6.4, and 7.5% acetonitrile) to resolve nucleotide mixtures loaded in sample buffer (25 mM HEPES, pH 7.4, 140 mM KCl, 15 mM NaCl). **Figure 4** illustrates the excellent nucleotide resolution observed for a mixture of GMP, GDP, GTP, GMPPNP, and GMPPN nucleotide standards (40 µM each), where peak assignments were determined by separate HPLC analyses of the individual nucleotides. As there is no substantive overlap of peaks it is straightforward to quantitate each nucleotide by integrating its peak area and comparing to a standard curve generated for nucleotide samples of known concentration. The order that nucleotides are eluted from the reverse-phase column are in line with what other studies have identified, yielding the elution times summarized in **Table 1** [44, 54]. GMPPN, which to our knowledge has not been run before, is found to elute between GMP and GDP (Fig 4, Table 1). For unknown reasons, GMP consistently displays two peaks as seen previously [44].

**Figure 3.**
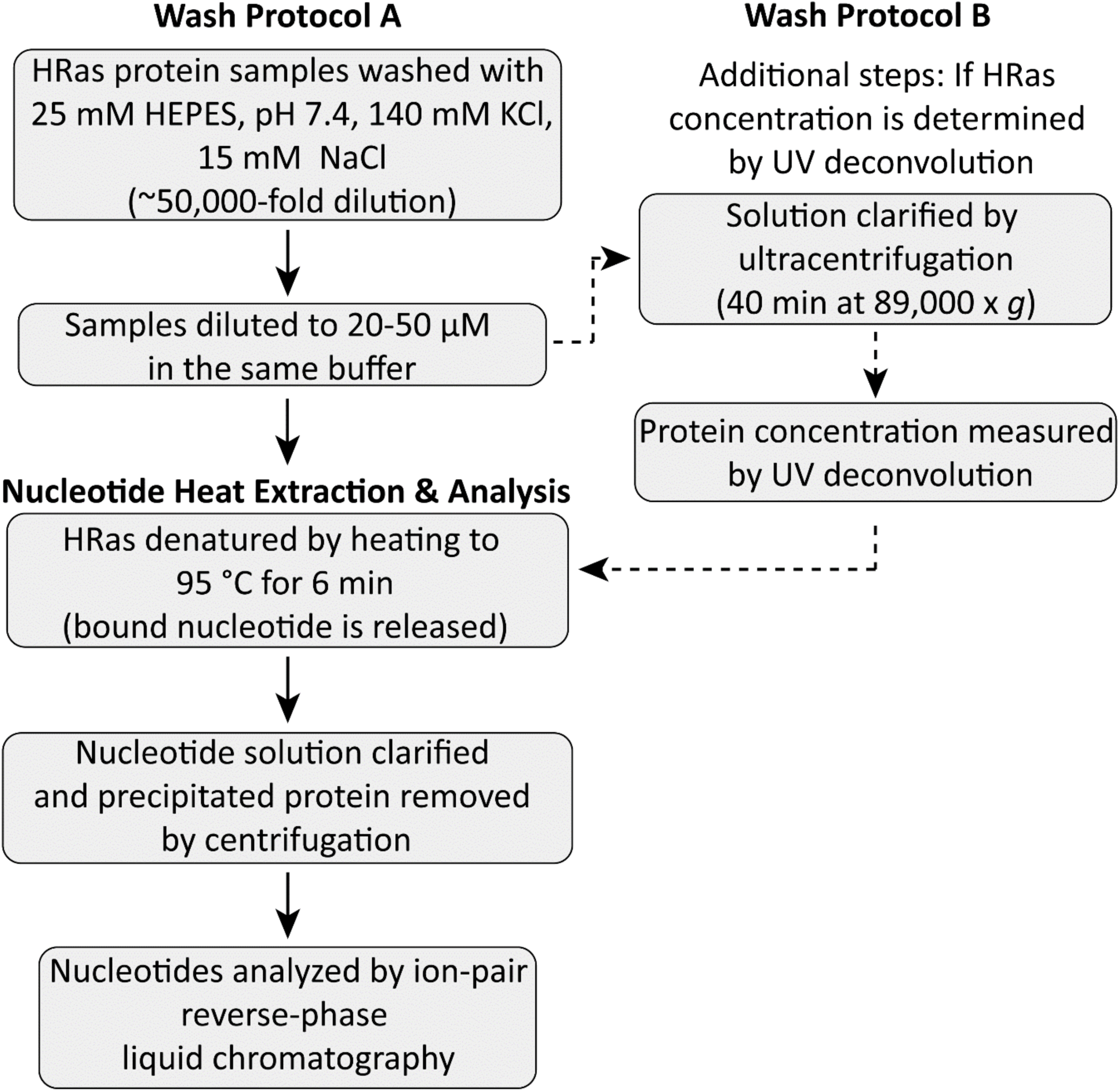
Protocols employed to wash HRas proteins and isolate their nucleotide mixtures for HPLC analysis. Flow chart summarizing the two nucleotide wash protocols employed herein. Each protocol begins with HRas previously loaded with the desired nucleotide. Wash Protocol A: HRas is washed extensively to remove unbound nucleotides, then a heating step denatures the protein and extracts its bound nucleotides, and finally denatured protein is removed by centrifugation to yield the extracted nucleotides analyzed by HPLC. Wash Protocol B: Same as A, with additional steps to prepare the sample both for nucleotide analysis via HPLC and protein analysis via UV deconvolution [34].

**Figure 4.**
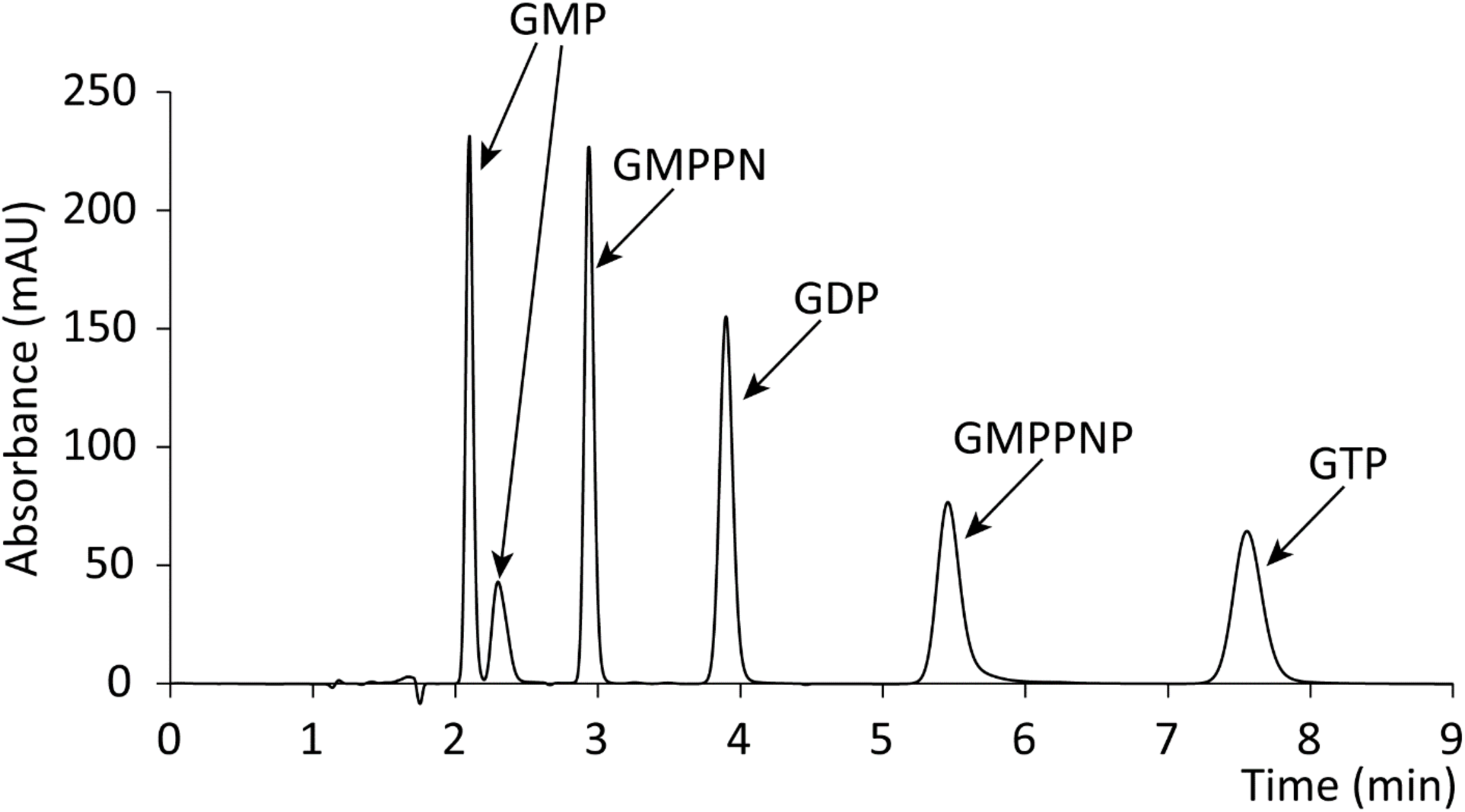
Control HPLC-UV chromatogram of guanine nucleotide standards. Indicated is a representative chromatogram obtained for a mixture of nucleotide standards containing 40 µM each of GMP, GMPPN, GDP, GMPPNP and GTP. Two peaks are observed for GMP, with retention times of 2.1 and 2.3 min, respectively, while the remaining nucleotides display retention time ranges as follows: GMPPN 2.9-3.0 minutes; GDP 3.8-4.0 minutes; GMPPNP 5.1-5.6 minutes; and GTP 6.8-7.8 minutes.

**Table 1.** Retention times for guanosine and the guanine nucleotides GMP, GMPPN, GDP, GMPPNP and GTP. The indicated retention time ranges were defined by HPLC measurements over a 1+ year period using two identical Reverse-phase C18 columns.

| **Nucleotide** | **Retention time**<br>**(mins)** |
| --- | --- |
| Guanosine | 1.8 |
| GMP: Major peak | 2.1 |
| Minor peak | 2.3 |
| GMPPN | 2.9-3.0 |
| GDP | 3.8-4.0 |
| GMPPNP | 5.1-5.6 |
| GTP | 6.8-7.8 |

### 3.2 Effect of Heat on Guanine Nucleotides

To isolate the nucleotide mixture bound to HRas for HPLC, we used a heat treatment step to denature the protein and release its bound nucleotides. Optimization of the heat treatment was carried out to facilitate quantification of the bound nucleotide mixture, including careful measurements of the effects of heating on the three major guanine nucleotides bound to HRas in our studies: GMPPNP, GTP and GDP. A number of previous studies have also utilized thermal denaturation of small GTPases, followed by clarification of the analyte solution prior to qualitative or quantitative analysis of nucleotide-loading by HPLC-UV, with heating times ranging from 2 to 5 min [48, 52, 53]. Accordingly, we decided it was necessary to investigate the effect on heat, not just on GMPPNP, the chosen hydrolysis-resistant GTP analog for these studies, but also on GTP and GDP, the physiologic nucleotides which bind to the active site of HRas proteins with high affinity [14]. Each nucleotide stock was divided into identical aliquots which were subjected to one of the following conditions: no heat, 3 min of heating at 95 °C, or 6 min of heating at 95 °C, prior to analysis by HPLC-UV.

When unheated GMPPNP was analyzed by HPLC, two peaks were eluted. The major, intact GMPPNP peak accounted for 94 ± 1% of the total peak area, while the minor hydrolysis product GMPPN accounted for 6 ± 1% of the total area (**Figure 5A**, **Figure 6A** and **Supplemental Figure 2**). By contrast, heating GMPPNP at 95 °C for 3 min caused a dramatic shift in the fraction of intact nucleotide relative to the hydrolyzed form as measured by peak area. In this case, approximately 12% of the GMPPNP remained intact and 88% was hydrolyzed to GMPPN (**Figure 6A**). Finally, heating GMPPNP at 95 °C for 6 min was sufficient to bring the hydrolysis of GMPPNP nearly to completion, yielding approximately 1% GMPPNP and 99% GMPPN (**Figure 5B** and **Figure 6A** and **Supplemental Figure 2**).

**Figure 5.**
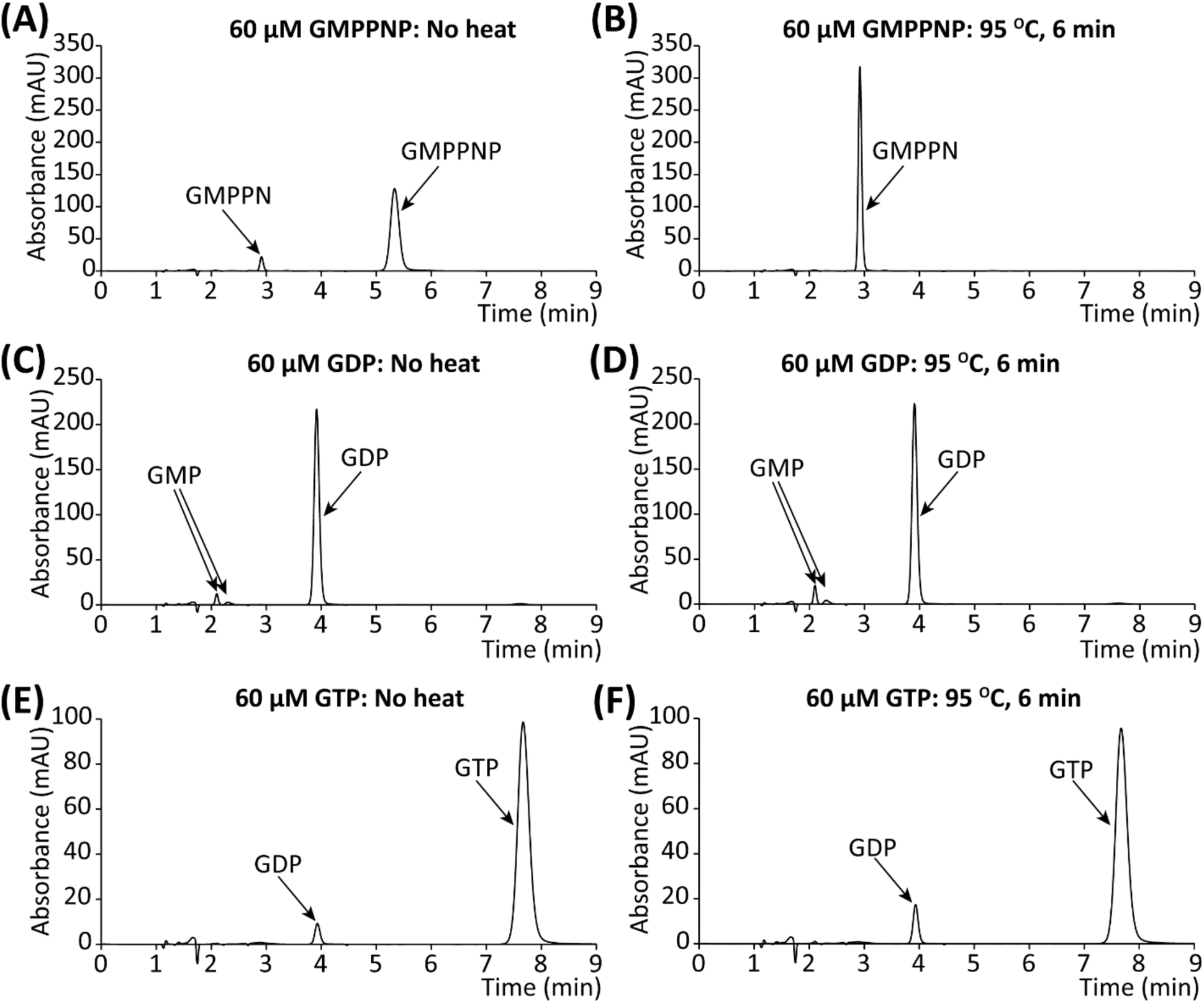
HPLC-UV chromatograms showing the effects of heat treatment on the guanine nucleotides GMPPNP, GDP and GTP. **(A)** Non-heat-treated 60 µM GMPPNP displays a minor peak (GMPPN) and a major peak (GMPPNP). (**B**) Heat-treated 60 µM GMPPNP (95 °C for 6 min) displays the same two peaks, but now the majority of GMPPNP has been hydrolyzed to GMPPN. (**C**) Non-heat-treated 60 µM GDP displays minor peaks (GMP) and a major peak (GDP). **(D)** Heat-treated 60 µM GDP (95 °C for 6 min) displays the same peaks with a small fraction of GDP, which remains the major component, converted to GMP. (**E**) Non-heat-treated 60 µM GTP displays a minor peak (GDP) and a major peak (GTP). **(F)** Heat-treated 60 µM GTP (95 °C for 6 min) displays the same peaks with minor conversion of GTP, which remains the major component, to GDP. For additional data and time courses see **Supplemental Figure 3**.

**Figure 6.**
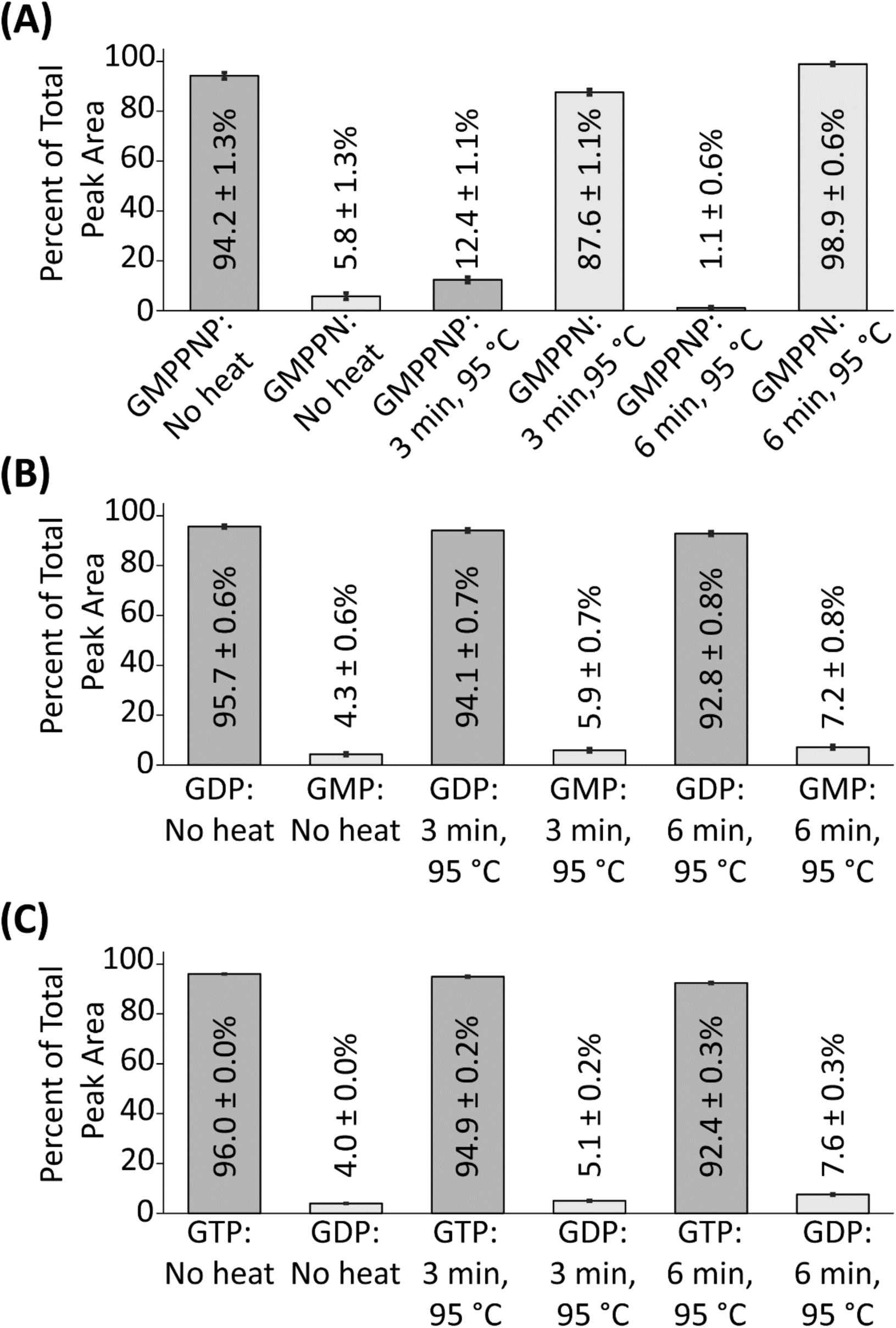
Quantified effects of heat treatment on the guanine nucleotides GMPPNP, GDP and GTP. Shown are the peak percentage areas observed for **(A)** GMPPNP and GMPPN when samples of 60 µM GMPPNP are subjected to no heating, 95 °C for 3 min, and 95 °C for 6 minutes; **(B)** GDP and GMP when samples of 60 µM GDP are subjected to the same three conditions; **(C)** GTP and GDP when samples of 60 µM GTP are subjected to the same three conditions.

When unheated GDP was analyzed by HPLC-UV, a single major peak corresponding to GDP accounted for 96% of the total peak area, while minor peaks corresponding to GMP accounted for 4% of the total peak area (**Figure 5C** and **Figure 6B**). When identical samples of GDP were subjected to heating at 95 °C for 3 or 6 min, only slight changes in the respective nucleotide species peak areas could be observed: the GDP peak area decreased to approximately 94% or 93%, respectively and the GMP total peak area increased to 6% or 7%, respectively (**Figure 6B**). Heating at 95 °C, therefore, appears to convert a small fraction of GDP to GMP over the timeframe used in these studies (**Figure 5D** and **Figure 6B**). As heating times were increased beyond 6 min up to 15 min, increased levels of GMP were generated indicating the importance of carefully controlling the heating time (**Supplemental Figure 3**).

When unheated GTP standard was analyzed by HPLC-UV, two peaks were observed: a major GTP peak accounting for 96% of the total peak area, and a minor GDP peak representing 4% of the total peak area (**Figure 5E** and **Figure 6C**). When an identically prepared GTP sample which had been subjected to heating at 95 °C for 3 or 6 min was analyzed by HPLC-UV, again two peaks could be observed with identical retention times to those of the unheated sample. In this case, the GTP peak accounted for 95% or 92% of the total peak area, respectively, while the GDP accounted for 5 or 8% of the total peak area, respectively (**Figure 5F** and **Figure 6C**). This trend of slightly increased levels of GTP hydrolysis as a consequence of heating was highly reproducible, both with identical 60 µM GTP stock solutions and also with other GTP stock solutions of different concentrations (data not shown). It was also observed that levels of GTP hydrolysis increased as heating times were increased beyond 6 minutes, (**Supplemental Figure 3**).

The findings indicated that a heat treatment of 95 °C for 6 min is well suited for quantifying the GMPPNP, GTP and/or GDP nucleotides released from HRas by thermal denaturation. This treatment yielded nearly complete conversion of GMPPNP to GMPPN, while retaining 96-97% of the starting GTP and GDP populations. Notably, GMPPNP bound to HRas is kinetically stable (the intrinsic GMPPNP hydrolysis rate of HRas = 25.6 X 10^-5^ min^-1^ at 37 °C, yielding an effective lifetime exceeding 2 days [28]). Moreover, the GMPPNP stock reagent used to load HRas is 94% pure (with 6% GMPPN also present), and a phosphatase inhibitor is used to block any contaminating phosphatase activity during nucleotide loading. Thus, the GMPPN bound to HRas prior to the heat extraction step is minimal. Additional studies confirmed that the heat extraction releases virtually all of the bound nucleotides from the protein, as evidenced by the measured equimolar ratio of total extracted nucleotides:total protein (**Section 3.4**). It follows that the GMPPN measured after heat extraction is a useful proxy, with an accuracy approaching 94%, for the GMPPNP bound to the starting HRas population.

### 3.3 Resolution and Identification of Nucleotides Bound to HRas and Mutant HRas Proteins

To resolve and identify the mixture of guanine nucleotides bound to HRas or one of its mutants, each purified protein was first washed extensively, then its bound nucleotides were heat-extracted and analyzed by HPLC. The extensive washing with sample buffer (net 50,000-fold dilution via buffer exchange with 25 mM HEPES, pH 7.4, 140 mM KCl, 15 mM NaCl) ensured removal of unbound nucleotides and any residual components of the protein preparation buffers that would interfere with HPLC analysis. Subsequent to buffer exchange, the protein was diluted to a standard concentration range (20-50 µM) and the nucleotide extraction was carried out by heating to 95 °C for 6 min, yielding protein denaturation and nucleotide release. As reported in **Section 3.2** above, the heating step has reproducible, quantified effects on the nucleotide population, generating efficient conversion of GMPPNP to GMPPN with minimal effects on GTP and GDP.

The resulting heat-extracted nucleotide mixtures were then subjected to HPLC-UV analysis. Each HPLC run also included a standard mixture of nucleotides to confirm normal nucleotide elution times and resolution. **Figure 7A-F** shows representative HPLC chromatograms obtained for six different HRas proteins loaded with GMPPNP during the protein purification process: standard HRas (the C118S/C181S mutant) and five HRas mutants (G12S, E37G, D38E, E63K and Y64G, each constructed in the standard HRas background). Similarly, **Figure 8A-B** shows representative chromatograms for standard HRas loaded with GMPPNP or GDP during protein purification, respectively. Each of the chromatograms in **Figures 7** and **8** yield excellent peak resolution and unambiguous identification of the major guanine nucleotide species that were bound in the active site of the washed protein. The retention times for the peaks observed in these chromatograms are entirely within the ranges observed for the single and mixed nucleotide standards of GMP, GMPPN, GDP, GMPPNP and GTP reported in **Figure 4** and **Table 1**.

**Figure 7.**
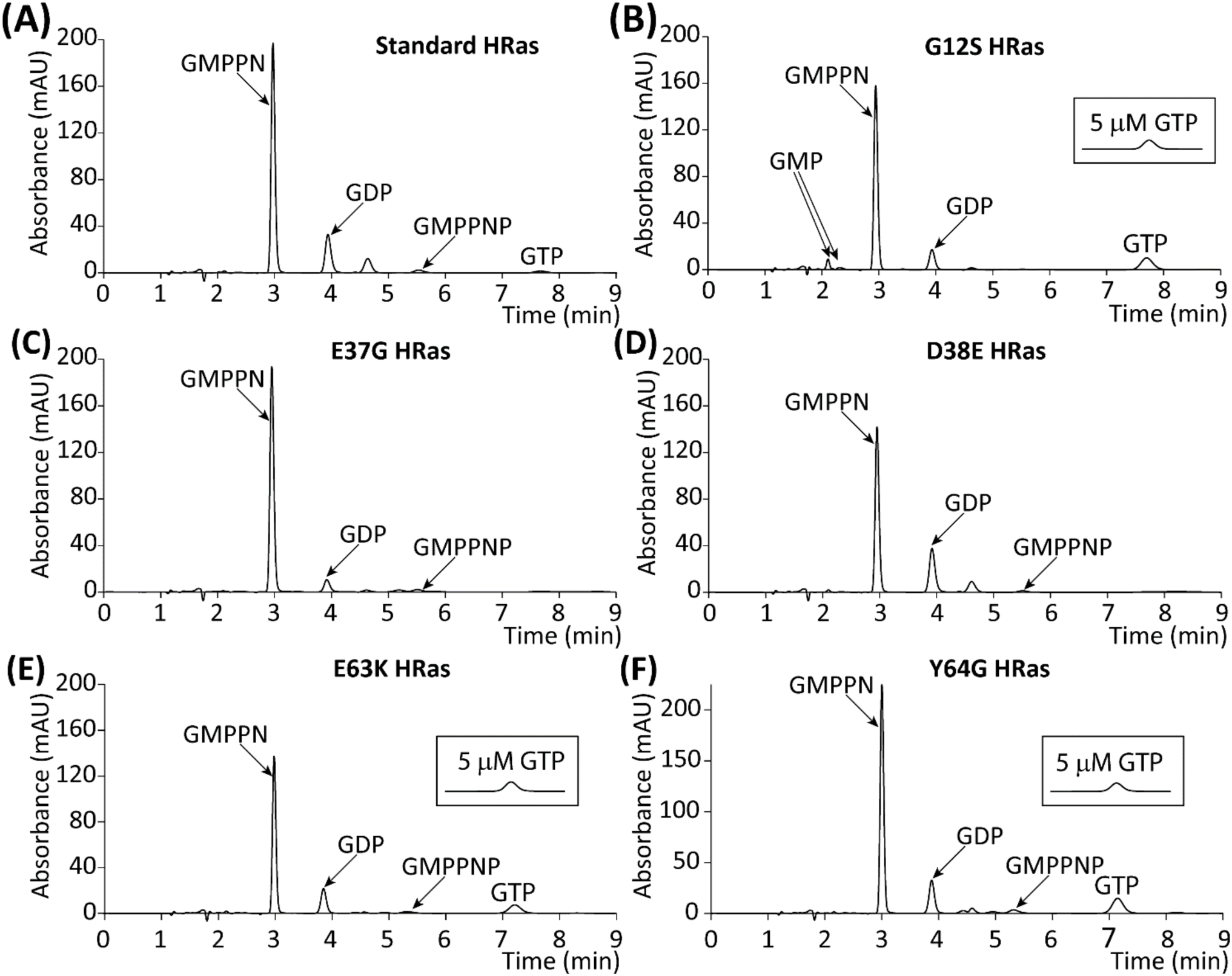
HPLC-UV resolution and identification of nucleotides isolated from HRas and HRas mutants loaded with GMPPNP. Each chromatogram shows the nucleotide mixture isolated from the indicated GMPPNP-loaded HRas protein by using Protocol A (Figure 3) to remove unbound nucleotides, then heat-extract the bound nucleotides. All proteins possessed the same engineered background mutations C118S and C181S, including **(A)** standard HRas and HRas mutants **(B)** G12S (known gain-of-function), **(C)** E37G, **(D)** D38E, **(E)** E63K, and **(F)** Y64G. Peaks corresponding to GMP, GMPPN, GDP, GMPPNP and GTP are indicated. For subsequent quantitation of peaks, a dilution series of standards was run in parallel to all samples. This series was created by heating equimolar GMPPNP and GDP standards (95 °C for 6 min) then diluting from 60 µM to 7.5 µM. For the G12S, E63K and Y64G mutants a separate dilution series was also run for GTP, created by heating GTP (95 °C for 6 min) then diluting from 10 µM to 1.25 µM (inset shows 5 µM). All data were processed in an identical manner (**Section 2.8**) yielding the quantified nucleotide percentages summarized in **Table 2** and **Supplemental Table 1**.

**Figure 8.**
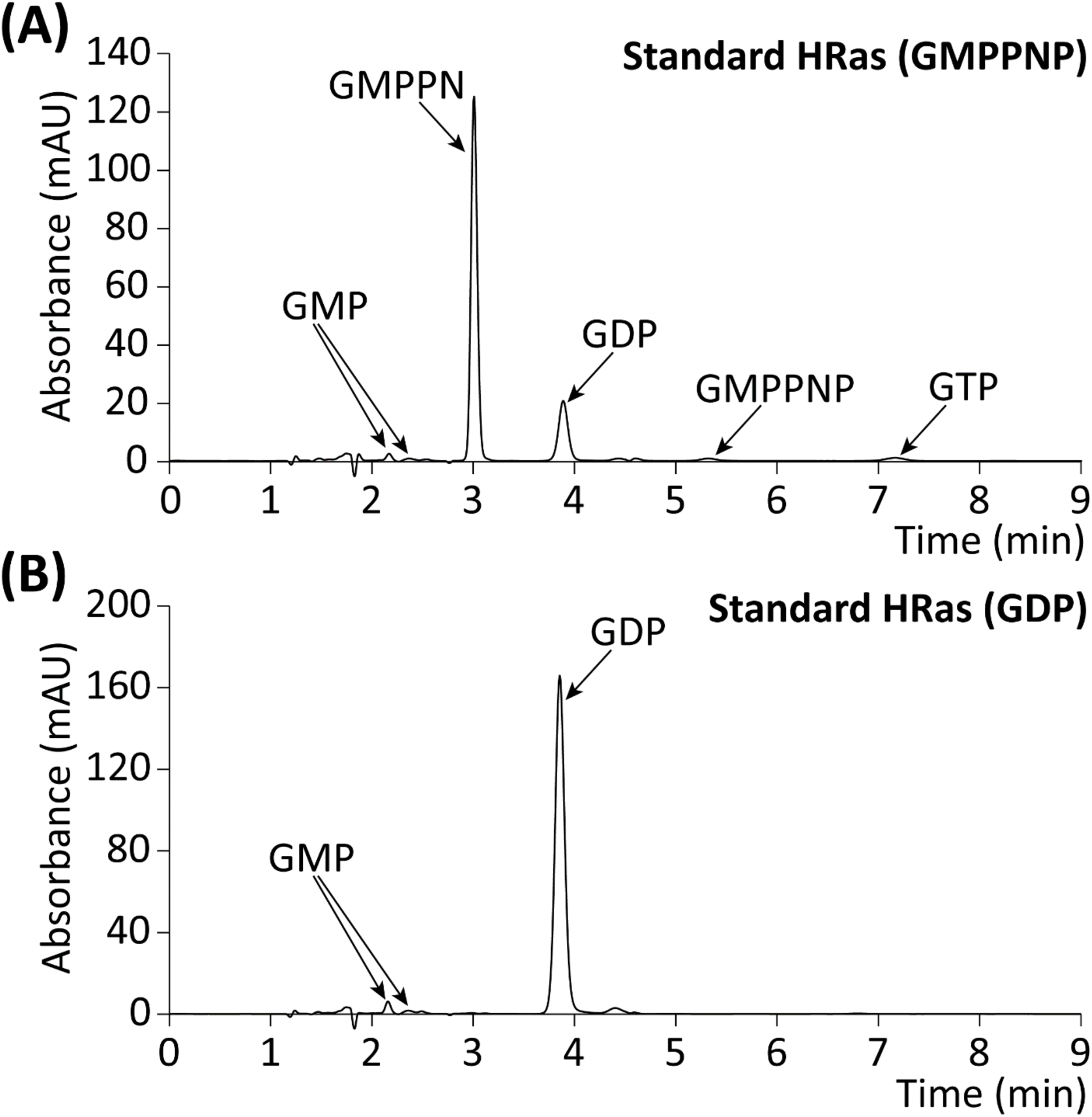
HPLC-UV resolution and identification of nucleotides isolated from HRas loaded with GMPPNP or GDP. Same as Figure 7 except that standard HRas was loaded with **(A)** GMPPNP or **(B)** GDP, then was subjected to Wash Protocol B to remove unbound nucleotides and heat extract the bound nucleotides (Figure 3). The use of Wash Protocol B enabled parallel quantitation of the HRas protein concentration via UV deconvolution [34].

**Table 2.** Quantities of identifiable guanine nucleotdies found in standard and mutant forms of HRas as determined by percentage peak area. ^¥^ Samples were generated using wash protocol A. In this case, a total of two samples were applied to the HPLC system, and measurements recorded in triplicate. * Samples were generated using wash protocol B. In this case, a total of three or six samples were applied to the HPLC system, and measurements recorded in triplicate.

| HRas | GMP (%) | GMPPN (%) | GDP (%) | GMPPNP (%) | GTP (%) | Nucleotides arising from active HRas (%) | Nucleotides arising from inactive HRas (%) | Range of Average [Total Nucleotide], (μM) | Range of Average [Total Protein], (μM) & (Mole Ratio of Nucleotide : Protein) |
| --- | --- | --- | --- | --- | --- | --- | --- | --- | --- |
| HRas*<br>(2 X 3) | 0.8 ± 0.7 | 75 ± 2 | 18.5 ± 0.5 | 3 ± 1 | 2.5 ± 0.7 | 81 ± 4 | 19 ± 1 | 48.1–48.2 | N/A |
| G12S*<br>(2 X 3) | 8 ± 2 | 70 ± 2 | 9 ± 2 | 0.4 ± 0.5 | 14.4 ± 0.3 | 83 ± 3 | 17 ± 4 | 39.6–40.2 | N/A |
| E37G*<br>(2 X 3) | Not measurable | 87.3 ± 0.5 | 7.4 ± 0.8 | 3 ± 1 | 2 ± 1 | 93 ± 3 | 7.4 ± 0.8 | 40.3–47.8 | N/A |
| D38E*<br>(2 X 3) | 1.7 ± 0.4 | 67.8 ± 0.7 | 27.4 ± 0.9 | 2.9 ± 0.2 | 0.3 ± 0.6 | 71 ± 2 | 29 ± 1 | 33.2–37.7 | N/A |
| E63K*<br>(2 X 3) | 1.3 ± 0.7 | 67 ± 1 | 16.5 ± 0.5 | 3 ± 1 | 11.9 ± 0.6 | 82 ± 3 | 18 ± 1 | 33.2–37.7 | N/A |
| Y64G*<br>(2 X 3) | 0.7 ± 0.4 | 67.8 ± 0.8 | 14.9 ± 0.2 | 2.5 ± 0.8 | 14.1 ± 0.2 | 84 ± 2 | 15.6 ± 0.6 | 49.4–57.1 | N/A |
| HRas<br>(6 X 3) | 2.8 ± 0.8 | 74 ± 2 | 19.0 ± 0.8 | 2.1 ± 0.7 | 2.8 ± 0.8 | 78 ± 3 | 22 ± 2 | 27.2–42.1 | 29.4–45.6<br>(1:1.04 ± 0.03) |
| HRas<br>GDP*<br>(3 X 3) | 3.9 ± 0.7 | N/A | 96.1 ± 0.7 | N/A | Not measurable | Not measurable | 100 ± 1 | 21.3–41.8 | 20.0–40.1<br>(1:0.95 ± 0.01) |

### 3.4 Nucleotide Quantitation and Reconstruction of the Bound Nucleotide Population

For each standard or mutant HRas protein, the HPLC chromatogram of its heat-extracted nucleotide population was used to reconstruct both (i) the mixture of nucleotides bound to the original protein prior to nucleotide heat extraction, and (ii) the fraction of the original protein occupied by an activating or inactivating nucleotide, respectively, as follows. First, the nucleotides present in the heat-extracted mixture were identified and quantified by assigning their characteristic elution times and integrating the A252 nm signals of their HPLC peaks. This nucleotide quantitation is facilitated by the spectral dominance of the guanine base, which ensures that all of the relevant nucleotides have the same ε_252_ =13.7×10^3^ M^-1^ cm^-1^ [40] Second, the resulting nucleotide integrals were converted to moles by interpolation or extrapolation on standard curves plotting the integrals obtained for increasing known quantities of the same nucleotides subjected to the standard heat treatment of 95 °C for 6 min, as illustrated in **Figure 9**. Third, the moles of different nucleotides were totaled, and the mole percent of each nucleotide was calculated. Fourth, the fraction of the original protein in the activated state was calculated by summing the mole percents of GMPPNP, GMPPN, and GTP, and the fraction of original protein in the inactive state was calculated by summing the mole percents of GDP and GMP. The latter calculations assume that a) the original mole percent of GMPPNP is the sum of the measured GMPPNP and GMPPN mole percents following heat extraction, and b) the original mole percent of GDP is the sum of the measured GDP and GMP mole percents following heat extraction. Finally, for the proteins subjected to a longer analysis procedure (Wash Protocol B in **Figure 3**) that included determination of the total protein concentration, the mole ratio of total bound nucleotide : total protein was determined.

**Figure 9.**
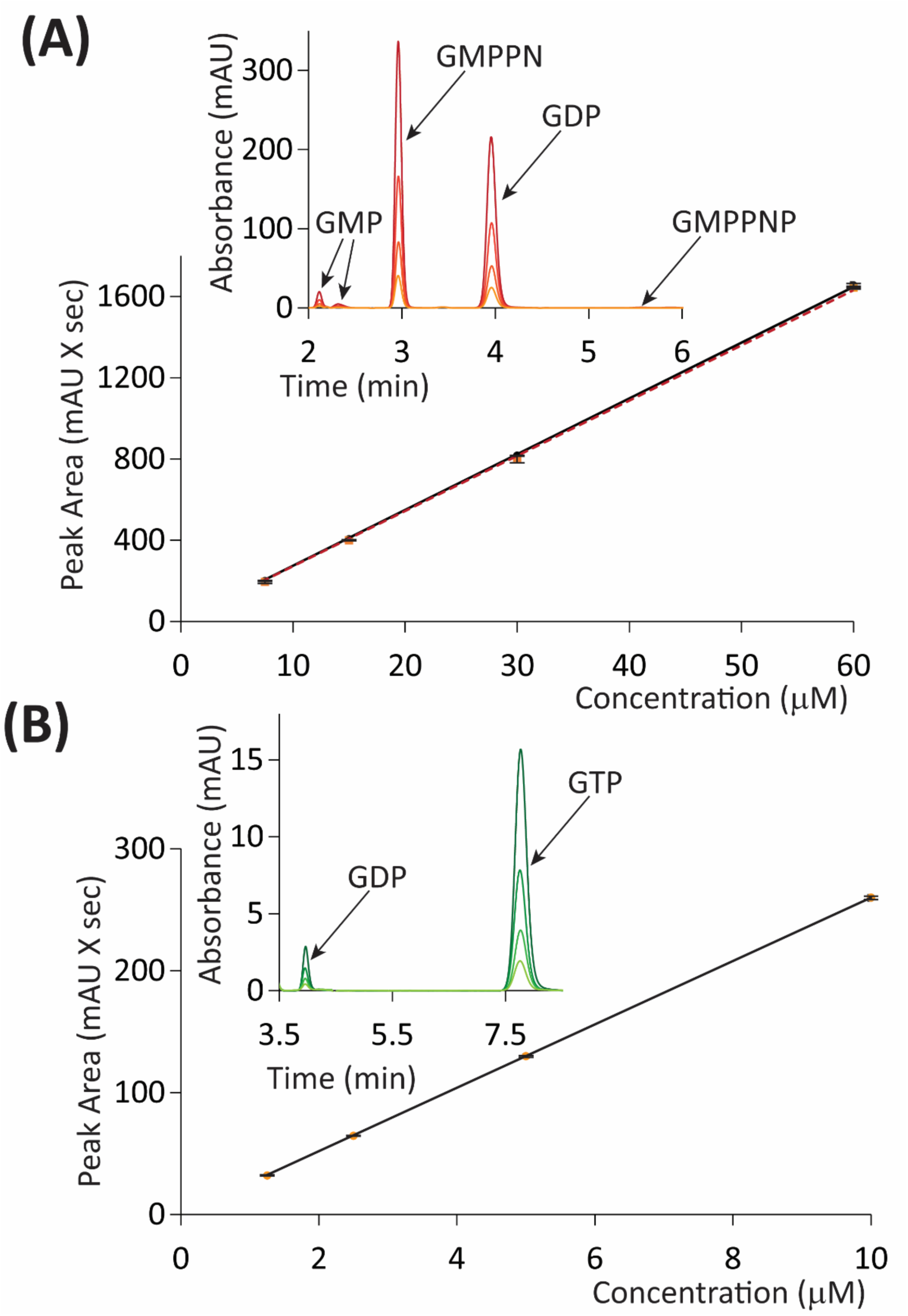
Representative standard curves used to quantify the concentrations of individual nucleotides in the nucleotide mixtures isolated from HRas proteins. **(A)** Shown are standard curves plotting peak area vs concentration for mixtures of GMPPNP (solid line) and GDP (dashed line) heated at 95 °C for 6 min, then used to create a dilution series of 60 µM, 30 µM, 15 µM, and 7.5 µM. Inset are overlaid HPLC-UV chromatograms of the resulting peaks. **(B)** Standard curve generated for GTP heated at 95 °C for 6 min, then diluted from 10 to 1.25 µM. For each standard curve, the indicated linear best fit yielded a slope that was used to quantify the concentration (concentration = peak area / slope) of the corresponding nucleotide in chromatograms of heat-extracted nucleotide mixtures.

**Table 2** summarizes the measured average parameters for each protein analyzed by HPLC in **Figure 7** and **Figure 8**. The findings show that in general, the mole fractions of the multiple nucleotide components extracted from a given HRas protein or mutant are highly reproducible between independent replicates measured on the same day or different days. In addition, for the washed HRas samples subjected to both protein [34] and nucleotide analysis, the mole ratio of nucleotide : protein was found to approach, within error, the known 1:1 stoichiometry. It follows that the HRas proteins retained their native, extremely high nucleotide affinities throughout their preparation and extensive washing, and that the washing steps were sufficient to remove virtually all unbound nucleotide.

The findings revealed two subsets of GMPPNP-loaded HRas proteins that differed in their levels of residual GTP that was loaded in the cell prior to isolation and purification. Three GMPPNP-loaded proteins, namely standard HRas and the E37G and D38E mutants, exhibited low levels of residual GTP. The HPLC chromatogram for heat-extracted nucleotide mixtures from standard, GMPPNP-loaded HRas was dominated by GMPPN (75 ± 2% of total guanine peak area) and GDP (18.5 ± 0.5% of the attributable guanine peak area). A smaller quantity of GMPPNP (3 ± 1% of the attributable guanine peak area) that survived the heat treatment was also observed, as were low levels of GTP (2.5 ± 0.7%), and GMP (0.8 ± 0.7%). The fraction of the starting, washed HRas population in the activated state prior to nucleotide extraction was operationally defined by totaling the GMPPNP, GMPPN, and GTP peaks, yielding 81 ± 4% of the total identifiable guanine nucleotide peak area (**Table 2**). Similarly, the fraction of the starting HRas population in the inactive state was operationally defined by adding the GDP and GMP peaks, yielding 19 ± 1% of the total attributable guanine nucleotide peak area. Comparable nucleotide profiles were observed for the HRas mutants E37G (93 ± 3% activated prior to nucleotide extraction) and D38E (71 ± 2% activated prior to nucleotide extraction). In each case, GMPPN and GDP were the major nucleotide species present, with minor components of GMPPNP, GMP and GTP (**Figure 7** and **Table 2**).

Three other GMPPNP-loaded mutant proteins, namely the extensively studied ‘gain-of-function’ G12S as well as newly studied E63K and Y64G, retained higher levels of residual GTP that survived cell isolation, purification, GMPPNP loading, washing, and nucleotide heat-extraction. For G12S, GMPPN was still the major guanine nucleotide present (70 ± 2% of the attributable guanine nucleotide peak area), while GTP was also present in significant quantities (14.4 ± 0.3% of the total guanine nucleotide peak area). In this case, GDP comprised the third largest component (9 ± 2% of the guanine nucleotide peak area) while GMP and GMPPNP were also present in lower concentrations. Prior to nucleotide extraction, the fractional populations of activated and inactivated G12S protein were estimated to be 83 ± 3% and 17 ± 5%, respectively. Similarly, the E63K and Y64G HRas mutants retained significant levels of residual GTP (11.9 ± 0.6% and 14.1 ± 0.2%, respectively) and were activated to comparable levels prior to nucleotide extraction (82 ± 3% and 84 ± 2%, respectively). Notably, while G12S was previously reported to inhibit the HRas intrinsic GTPase activity, the present findings provide the first evidence that the disease-linked mutation E63K, and the designed Y64G also inhibit the intrinsic HRas GTPase.

As controls, **Figure 8** and **Table 2** compare the HPLC analyses of GMPPNP-loaded HRas and GDP-loaded HRas. These controls confirm the ability of the HPLC analysis to identify both highly activated and highly inactivated HRas populations. On average, the GMPPNP-loaded HRas analyses yielded a low GDP level of 22 ± 2% of the total guanine nucleotides and indicated that the net HRas activation prior to nucleotide extraction was 78 ± 3%. In contrast, GDP-loaded HRas yielded a high GDP level of 96.1 ± 0.7% of the total guanine nucleotides, with no measurable activation prior to nucleotide extraction (GMP was the only other measurable nucleotide on this occasion (**Figure 8** and **Table 2**)).

The reproducibility of the HPLC method is illustrated by **Supplemental Table 1**, which reports all of the independent sample measurements that contribute to the averages presented in **Table 2**. Notably, independent sample measurements are found to yield nucleotide compositions and activation parameters generally within error of each other indicating that the HPLC analysis is highly reproducible. In short, the extensive data presented herein indicates that the HPLC method is both reproducible and capable of distinguishing the subtle (**Figure 7**) and major (**Figure 8**) differences between the bound nucleotide populations of a wide array of HRas variants in different activation states.

## 4. DISCUSSION

Overall, the findings indicate that the new HPLC-UV method presented herein is highly reproducible and provides resolution, identification, and quantitation of the multiple nucleotides bound to a given HRas population. The resulting analysis of the bound nucleotide mixture enables quantitation of the fraction of the HRas population stabilized in the on-state by occupancy with an activating guanine nucleotide (GMPPNP or GTP in the present study), as well as the complementary fraction of the population stabilized in the off-state by occupancy with inactivating nucleotide (GDP). The method is illustrated by successful application to multiple standard HRas preparations and five HRas mutants loaded with the activating nucleotide GMPPNP, and to HRas loaded with the inactivating nucleotide GDP.

The new HPLC-UV method includes useful features adapted or modified from previous HPLC-UV methods used to examine the nucleotide loading, or the intrinsic GTPase activity of small GTPase proteins [41–53]. The majority of these studies utilize a reverse-phase column fitted with a pre-column and an isocratic or gradient elution profile where the mobile phase consists of phosphate buffer containing an ion-pairing agent such as tetrabutylammonium bromide, and a small percentage of acetonitrile. This solvent system partially denatures G proteins, yielding at least partial release of their bound nucleotide population. Often, the resulting samples are applied directly to the HPLC system, and a pre-column is used to trap the denatured G protein while passing the free nucleotide. While such an approach can be useful, it limits pre-column and column longevity. Thus, a number of alternative approaches have been developed to extract the tightly bound guanine nucleotides from the G protein by more fully denaturing the nucleotide-protein complex, followed by removal of the precipitated protein by centrifugation prior to HPLC-UV analysis. The two most commonly employed methods of protein denaturation involve either heating, or alternatively, incubating the protein with an acidic reagent, such as perchloric acid [48–53]).

Development of the present method began with the choice of GMPPNP and GDP as the nucleotides employed in functional studies to stabilize the HRas on- and off-states, respectively, together with the choice of heating to fully denature the G protein and quantitatively release its bound guanine nucleotides. GMPPNP, GTP-γ-S, and GppCH_2_p are commonly employed as hydrolysis-resistant GTP analogs since Ras proteins enzymatically hydrolyze these analogs up to 190-fold more slowly than GTP, enabling stabilization of the active conformation for subsequent functional and structural analyses [28]. Indeed, numerous three-dimensional structures of wild-type or mutant Ras proteins bound to GMPPNP, GTP-γ-S or GppCH_2_p have been deposited in the RCSB Protein Data Bank [11, 55]. We chose GMPPNP to stabilize the on-state due to its superior combination of properties as follows. (i) GMPPNP is readily commercially available at a high level of purity (94 ± 1% based on our HPLC-UV analysis of standard samples). Thus, recombinant Ras protein can be loaded with GMPPNP to trap a large fraction of the G protein population in the active state. (ii) A structure of a GTP:HRas complex has been derived from a crystal of caged-GTP:HRas, after photolytic cleavage of the cage group [8]. This structure confirmed that GMPPNP is an excellent surrogate of GTP in the HRas active site, with regards to the positioning of the nucleotide phosphate groups within the active site. (iii) Phosphatase inhibitors are commercially readily available and can be added at any stage of GMPPNP loading to fully block hydrolysis by low levels of *E. coli* alkaline phosphatase sometimes observed in recombinant Ras preps. (iv) GMPPNP is only very slowly hydrolyzed by HRas (k = 25.5 X 10^-5^ min^-1^ at 37 °C, approximately 110-fold more slowly than GTP hydrolysis, yielding an effective on-state lifetime exceeding 2 days. By contrast, Ras hydrolyzes GTP-γ-S 10-fold more rapidly than it hydrolyzes GMPPNP [28]. (iv) Most importantly for the present HPLC-UV method, upon heating GMPPNP hydrolyzes to GMPPN, which can be easily distinguished by HPLC-UV analysis from the other relevant nucleotides (GTP, GDP) and their hydrolysis products (GDP, GMP). As a result, GMPPN generated by heat extraction of nucleotides from the Ras protein can be quantified by HPLC-UV and tracked back to the starting, bound GMPPNP prior to heating. In contrast, GTP-γ-S and GppCH_2_p are hydrolyzed primarily to GDP and GMP respectively, which cannot be tracked back to the starting nucleotide [56]. Turning to functional studies of the inactive state, we chose GDP as the stabilizing nucleotide since it is the native ligand of the off-state, is highly resistant to further hydrolysis by the G protein or heat and is readily quantifiable by HPLC-UV.

Development of the method also included optimization of the heat extraction approach to isolate the bound nucleotides bound from HRas and prepare them for HPLC-UV analysis. Heat extraction had been previously employed in HPLC-UV studies [53]. However, we found that further development of the heat extraction method was needed to fully denature the HRas protein and extract the bound nucleotides. We also added post-heating centrifugation and filtration steps to virtually eliminate the denatured, precipitated protein, thereby yielding a clarified nucleotide mixture ideally suited for HPLC-UV analysis. We found that the optimized heating heating protocol (95 °C for 6 min) denatured virtually all of the HRas protein, while releasing the bound nucleotide population (see validation below). Notably, the present study also quantified the effects of the optimized heating protocol on each of the relevant nucleotides. Different nucleotides exhibited unique, highly reproducible extents of hydrolysis during the standard heat treatment used for extracting nucleotides from HRas (**Figure 6**). Thus, for the nucleotides relevant to our HRas functional studies, the heating protocol (95 °C for 6 min) yielded almost complete hydrolysis of GMPPNP to GPMPN (98.9 ± 0.6%); but only very limited hydrolysis of GTP to GDP (3.7 ± 0.3%), and of GDP to GMP (3 ± 1%). These findings provide the information needed for rigorous back-calculation of the starting bound nucleotide composition of the HRas population (prior to heating) from the HPLC-determined nucleotide composition of the heat-extracted nucleotide mixture.

Following isolation of the nucleotide mixture bound to HRas via heat extraction, the present method employs HPLC-UV analysis to resolve, identify and quantify the components of the nucleotide mixture. The procedure utilizes an analytical C18 reverse-phase column fitted with a pre-filter guard column. Nucleotide mixtures are loaded in sample buffer (25 mM HEPES, pH 7.4, 140 mM KCl, 15 mM NaCl) and run in an ionic, mixed-solvent mobile phase (92.5 mM KH_2_PO_4_, 9.25 mM tetrabutylammonium bromide, pH 6.4, and 7.5% acetonitrile). The approach provides excellent resolution of the relevant nucleotides (**Figure 4**), and the lack of overlap between peaks ensures straightforward quantitation of each component and its concentration by integration of its HPLC absorbance peak, followed by interpolation or extrapolation of its concentration on a standard curve generated by HPLC analysis of samples of the same component at known concentrations.

Application of the new HPLC-UV method to a total of four different preparations of standard and five mutant HRas proteins illustrated the reproducibility of the method and highlighted its ability to quantitatively analyze the bound nucleotide composition and signaling state of a given HRas sample. For each type of protein, **Table 2** summarizes the global averages of its nucleotide composition and signaling states measured over multiple samples, while **Supplemental Table 1** provides the numbers for individual samples. Comparison of multiple HPLC-UV analyses of samples from the same protein preparation reveal that the quantification of an individual nucleotide was typically reproducible to the ∼95% level for the major nucleotide(s), and to the ∼90% level for the minor nucleotide(s) (**Table 2** and **Supplemental Table 1**).

The known 1:1 stoichiometry of guanine nucleotide per protein molecule enabled a quantitative test of the overall validity of the new HPLC-UV method. Three independent measurements of the total extracted nucleotide concentration and the total protein concentration were carried out, each in triplicate, for standard HRas loaded with GMPPNP or GDP then washed. The excellent agreement observed between the total nucleotide concentration measured by HPLC-UV and the total G protein concentration measured by the independent G protein UV deconvolution method [34] provides strong validation of the HPLC-UV method, and confirms the virtually complete extraction (within 5%) of bound nucleotide from HRas provided by the optimized heat extraction procedure (**Table 2**).

Closer examination of the HPLC-UV data for the standard HRas and the mutants E37G and D38E enables quantitative analysis of their signaling state, which is crucial for rigorous HRas functional studies. Our standard HRas construct exhibits wild-type HRas activity in functional assays and serves as the background in which point mutations are introduced (see Methods and [35]). Loading standard HRas with GMPPNP, followed by extensive washing to remove unbound nucleotides, was found to yield an HRas population occupied by 78 ± 3% bound GMPPNP (the sum of the heat-extracted components GMPPNP and GMPPN), 18 ± 1% GDP, and 3 ± 1% GTP (the latter two nucleotides are residual from cellular expression). It follows that 81 ± 4% of the standard HRas was stabilized in the on-state by occupancy with an activating nucleotide (the sum of the fractional loadings with GMPPNP and GTP). By contrast, GDP-loaded standard HRas exhibited 100 ± 1% occupancy with GDP. Now that these GMPPNP/GTP and GDP-loaded standard HRas preparations have been quantified with respect to their nucleotide loading, they can be used in rigorous functional studies to ascertain the specific activities of the HRas on- and off-states, respectively. In contrast, when the mutant proteins E37G and D38DE were loaded with GMPPNP and washed, they exhibited fractional occupancies with the GMPPNP/GTP activating nucleotides of 93 ± 3% and 71 ± 2%, respectively. It follows that different HRas preparations and/or mutants exhibit significant variations in nucleotide loading, emphasizing the importance of HPLC-UV analysis of HRas samples employed in functional studies to quantify their fractional loading with activating and/or inactivating nucleotides, and thus their effective signaling state.

Analysis of the HRas mutants E63K, and Y64G reveal an unexpected pattern that suggests the existence of a previously unknown molecular disease mechanism for disease-linked mutations at these positions. Nearly all sidechain substitutions at the G12 and Q61 hotspot positions are linked to human cancers or other disease states, at least in part because mutations at these hotspot positions, including G12S, generally inhibit the intrinsic GTPase activity of HRas [22, 23, 57]. Consistent with this picture, the present findings show that GMPPNP-loaded G12S mutant possesses greater occupancy with residual GTP from cellular expression (14 ± 1%) than the GMPPNP-loaded standard HRas, E37G and D38E proteins (2.5 ± 0.7%, 2 ± 1%, and 0.3 ± 0.6 %, respectively) (**Table 2**). Surprisingly, like G12S (**Table 2**) and other G12 and Q61 hotspot mutants, the E63K and Y64G mutants exhibit high levels of residual GTP occupancy (11.9 ± 0.6% and 14.1 ± 0.2%, respectively) [22, 23, 57]. Moreover, Q61, E63 and Y64 all lie on Switch II in close proximity to each other and to G12 (within 5-7 Å) [9]. Based on these strong patterns, we hypothesize that disease-linked mutations at positions E63 and Y64, like disease-linked mutations at G12 and Q61, share a common molecular disease mechanism in which the mutation triggers inhibition of intrinsic GTPase activity and overpopulation of the activated, GTP-bound state. Mutations at the G12 and Q61 positions also block GTPase stimulation by GTPase activating proteins (GAPs), so it is possible that mutations at E63 and Y64 may also block GAP-stimulated GTPase activity. In short, we propose that some or all mutations at the E63 and Y64 positions disrupt intrinsic GTPase activity, and perhaps GAP regulation as well, thereby leading to excessive, constitutive Ras activation by bound GTP. The present results directly support this mechanism for the Ras E63K mutation, which has previously been identified as a heterozygous HRas mutation in patients with Costello syndrome, a rare congenital disorder affecting multiple organ systems, and is a human cancer-linked mutation in both KRas and NRas (Catalogue of Somatic Mutations in Cancer (COSMIC) database [20, 58]). Similarly, while HRas Y64G is an engineered mutant known to block interactions of Ras with its PI3K effector [29, 59] and is not currently known to be disease-linked, Y64H is a cancer-linked mutation observed in both HRas and KRas (COSMIC). This picture predicts that other mutations at the Y64 position may eventually be discovered as disease-linked mutations as well.

In summary, we have developed and optimized a reproducible method for resolving, identifying and quantifying the guanine nucleotides bound to HRas complexes. This method employs heat extraction of the bound nucleotides followed by ion-paired reverse-phase HPLC-UV analysis. The method is especially applicable to quantification of the fractional activation of HRas loaded with the GTP analog GMPPNP to stabilize the on-state, or the fractional inactivation of HRas loaded with GDP to stabilize the off-state. Such quantification of the fractional activation and inactivation of HRas samples is crucial to rigorous functional studies of on-off switching. Validation is provided by the excellent agreement between the total nucleotide concentration measured by the method and the total HRas concentration measured by UV deconvolution [34], as predicted by the known 1:1 stoichiometry of the complex. We expect this new HPLC-UV method to be generalizable to other Ras isoforms and, more broadly, to members of the extended Ras superfamily of G proteins.

## Acknowledgements

Work was funded by a National Institutes of Health (NIH) Grant R01 GM063235 (to J.J.F.), by a NIH Molecular Biophysics Traineeship T32 GM065103 (to G.H.S.), by a Beckman Scholar Award (to N.J.C.), and by a Colorado Biological Sciences Initiative Scholar Award (to J.G.M.).

## Abbreviations

GTP: (guanosine-5’-triphosphate)
GDP: (guanosine-5’-diphosphate)
GMP: (guanosine-5’-monophosphate)
GMPPNP: (GMPPNP, guanosine-5’-[β,γ-imido]triphosphate)
GMPPNH2: (guanosine-5’-[β-amino]-diphosphate)
GXP: (mixture of guanine nucleotides)
HRas: (H isoform of human Ras protein)

---

**Supplemental Figure S1.**
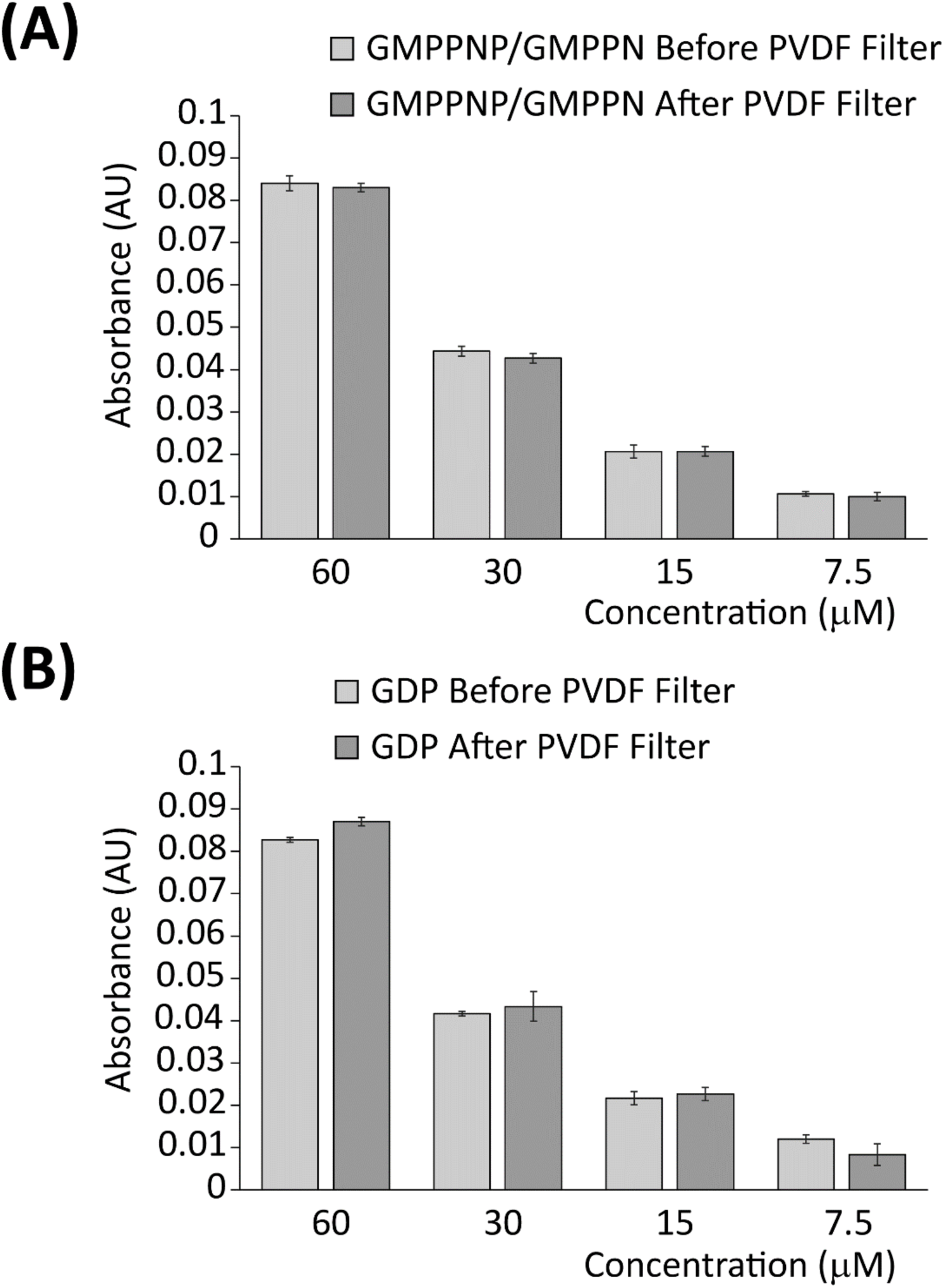
Nucleotide concentrations before and after PVDF filtration. Shown are average absorbance values recorded at a wavelength of 252 nm for **(A)** GMPPNP and **(B)** GDP that were heat-treated (95 °C for 6 min) then used to create a dilution series of 60 µM, 30 µM, 15 µM, and 7.5 µM before and after passing through a 0.20 µM PVDF filter. Data are averages of n = 3 replicates ± standard deviation.

**Supplemental Figure S2.**
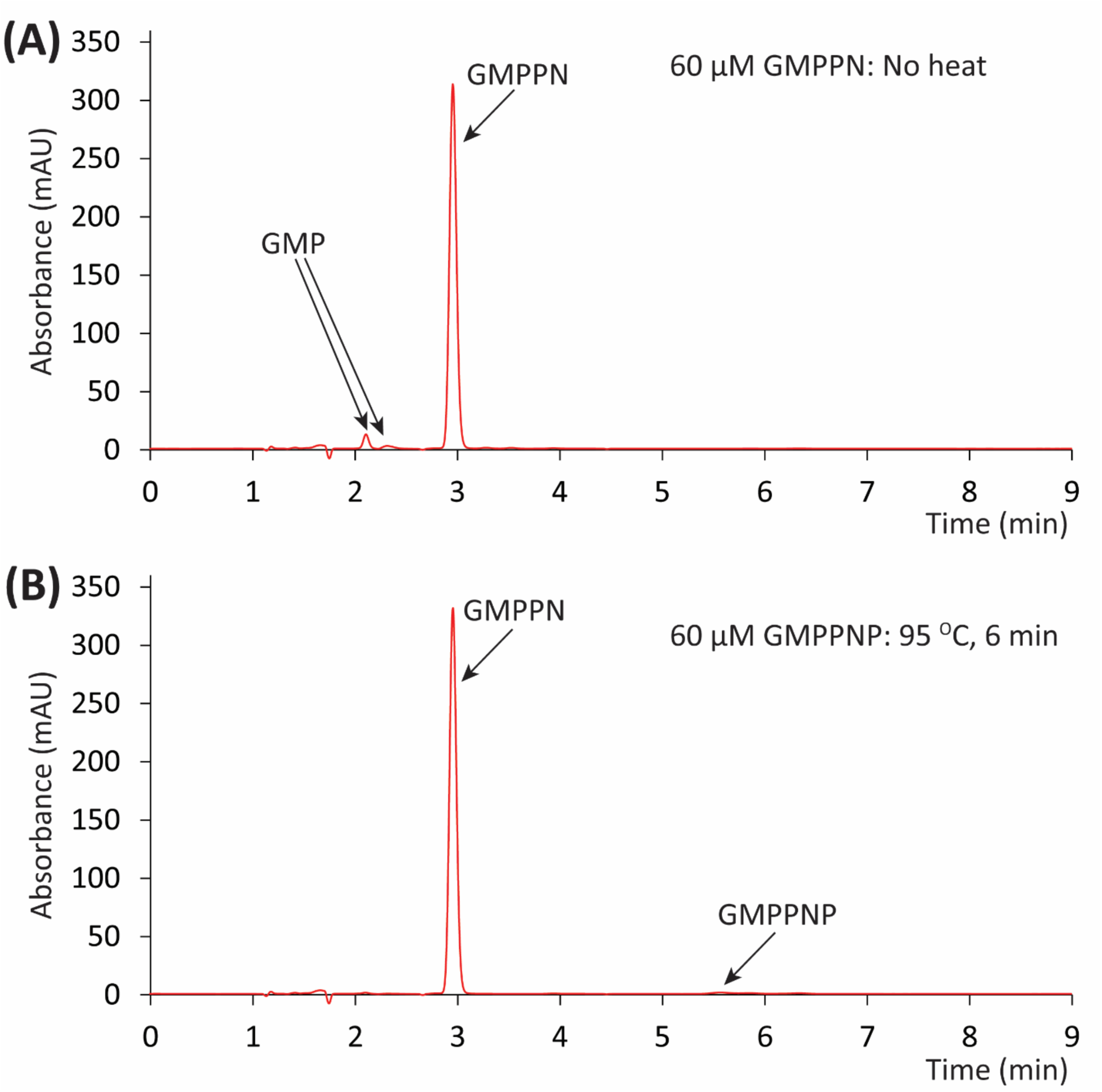
Control confirming that the major breakdown product of GMPPNP is GMPPN. **(A)** HPLC-UV chromatogram of commercially available, non-heat-treated GMPPN. Minor GMP peaks also indicated. (**B**) Heat-treated 60 µM GMPPMP (95 °C for 6 min), yielding a single peak that matches the retention time of the reference GMPPN.

**Supplemental Figure S3.**
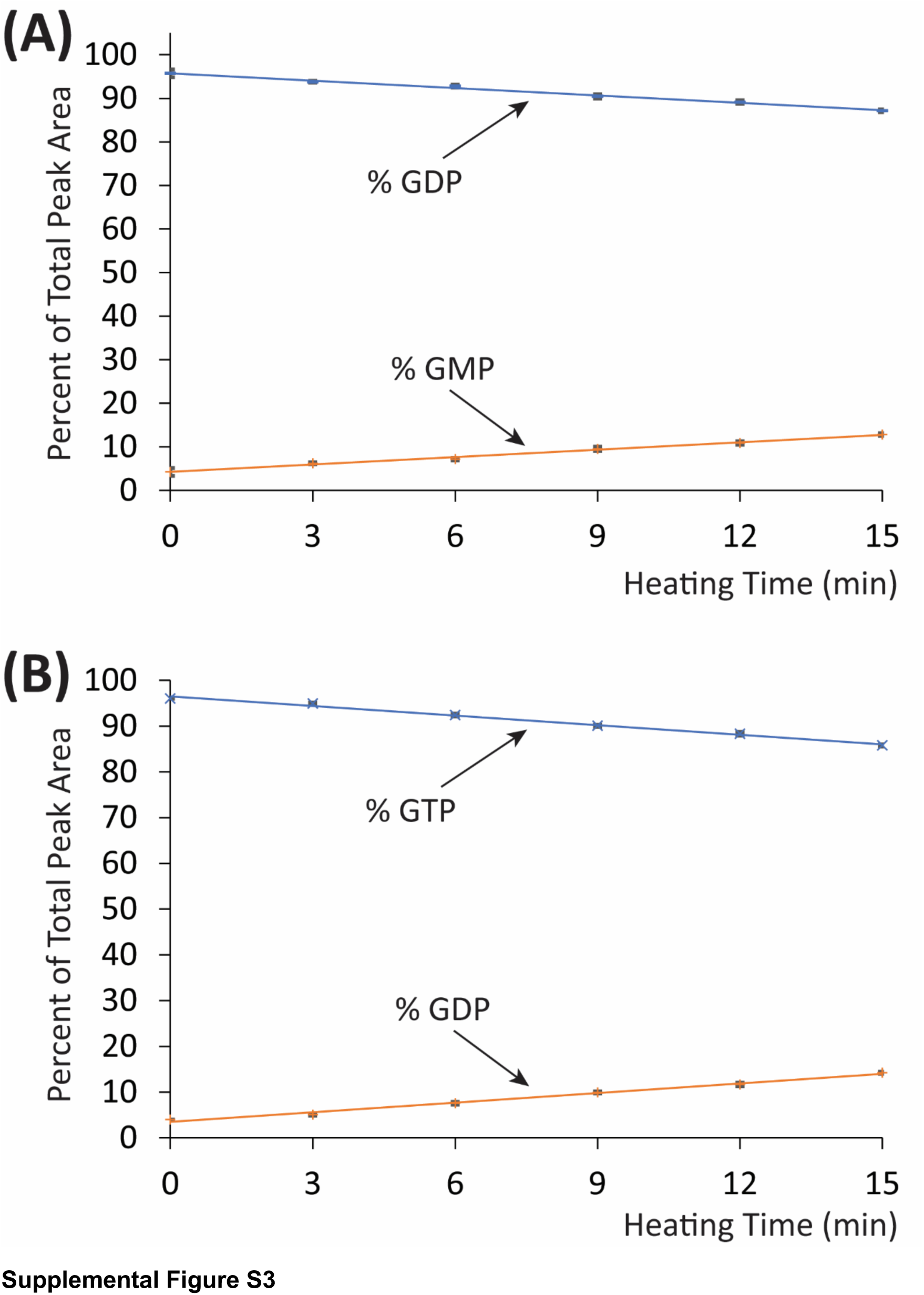
Time courses of heating effects on GDP and GTP. **(A)** GDP samples were placed in heat (95 °C) at t = 0 then sampled at the indicated 3-minute increments. As heating time increases GDP (blue) is hydrolyzed to GMP (orange). **(B)** GTP samples were placed in heat (95 °C) at t = 0 then sampled at the indicated 3-minute increments. As heating time increases GTP (blue) is hydrolyzed to GDP (orange).

**Supplemental Table S1.** **Calculated nucleotide concentrations derived from individual samples of standard and mutant forms of HRas.** ^¥^ Samples were generated using Wash Protocol A. * Samples were generated using Wash Protocol B. Each sample was measured in triplicate.

| HRas | [Protein], (μM)<br>&<br>(Nucleotide :<br>Protein<br>Stoichiometry) | Percentage<br>GMPNP/PN<br>(%) | [Total Nucleotide],<br>(μM) | Activating<br>Nucleotide :<br>Inactivating<br>Nucleotide<br>(%) : (%) | [Total<br>Activating<br>Guanine<br>Nucleotide],<br>(μM) | [Total<br>Inactivating<br>Guanine<br>Nucleotide],<br>(μM) |
| --- | --- | --- | --- | --- | --- | --- |
| HRas a<sup>‡</sup> | N/A | 77.6 ± 1.7 | 48.1 ± 2.0 | 80.2 : 19.8 | 38.6 ± 1.2 | 9.5 ± 0.8 |
| HRas b<sup>‡</sup> | N/A | 79.1 ± 0.9 | 48.3 ± 1.5 | 81.6 : 18.4 | 39.4 ± 1.0 | 8.9 ± 0.5 |
| G12S a<sup>‡</sup> | N/A | 67.3 ± 0.4 | 40.2 ± 1.1 | 82.1 : 17.9 | 33.0 ± 0.2 | 7.2 ± 0.9 |
| G12S b<sup>‡</sup> | N/A | 67.8 ± 2.2 | 39.6 ± 2.0 | 83.1 : 16.9 | 32.9 ± 0.9 | 6.7 ± 1.1 |
| E37G a<sup>‡</sup> | N/A | 91.0 ± 1.9 | 40.3 ± 1.9 | 92.3 : 7.7 | 37.2 ± 1.4 | 3.1 ± 0.5 |
| E37G b<sup>‡</sup> | N/A | 89.6 ± 0.9 | 47.8 ± 0.7 | 93.1 : 6.9 | 44.5 ± 0.6 | 3.3 ± 0.1 |
| D38E a<sup>‡</sup> | N/A | 70.9 ± 0.2 | 37.7 ± 0.8 | 71.4 : 28.6 | 26.9 ± 0.4 | 10.8 ± 0.4 |
| D38E b<sup>‡</sup> | N/A | 71.3 ± 0.9 | 33.2 ± 0.5 | 71.4 : 28.6 | 23.7 ± 0.3 | 9.5 ± 0.2 |
| E63K a<sup>‡</sup> | N/A | 69.6 ± 1.8 | 33.8 ± 1.2 | 82.2 : 17.8 | 27.8 ± 0.9 | 6.0 ± 0.3 |
| E63K b<sup>‡</sup> | N/A | 69.6 ± 1.8 | 34.9 ± 1.4 | 81.7 : 18.3 | 28.5 ± 0.8 | 6.4 ± 0.6 |
| Y64G a<sup>‡</sup> | N/A | 69.7 ± 0.3 | 57.0 ± 0.4 | 84.6 : 15.4 | 48.2 ± 0.2 | 8.8 ± 0.2 |
| Y64G b<sup>‡</sup> | N/A | 70.1 ± 1.4 | 49.5 ± 1.0 | 84.6 : 15.4 | 41.9 ± 0.7 | 7.6 ± 0.3 |
| HRas 1a<sup>*</sup> | 32.6 ± 0.1<br>(1.00 : 1.01) | 75.1 ± 1.0 | 32.2 ± 1.1 | 78.0 : 22.0 | 25.1 ± 0.7 | 7.1 ± 0.4 |
| HRas 1b<sup>*</sup> | 29.4 ± 0.3<br>(1.00 : 1.08) | 76.5 ± 1.1 | 27.3 ± 0.7 | 79.1 : 20.9 | 21.6 ± 0.6 | 5.7 ± 0.1 |
| HRas 1c<sup>*</sup> | 32.2 ± 0.2<br>(1.00 : 1.03) | 75.3 ± 1.0 | 31.4 ± 1.4 | 77.4 : 22.6 | 24.3 ± 0.7 | 7.1 ± 0.7 |
| HRas 2a<sup>*</sup> | 42.6 ± 0.5<br>(1.00 : 1.01) | 74.7 ± 0.9 | 42.0 ± 1.1 | 77.9 : 22.1 | 32.7 ± 0.5 | 9.3 ± 0.6 |
| HRas 2b<sup>*</sup> | 45.6 ± 0.5<br>(1.00 : 1.08) | 76.5 ± 0.5 | 42.1 ± 0.5 | 79.1 : 20.9 | 33.3 ± 0.3 | 8.8 ± 0.2 |
| HRas 2c<sup>*</sup> | 33.8 ± 0.1<br>(1.00 : 1.02) | 75.1 ± 1.4 | 33.1 ± 1.0 | 77.0 : 23.0 | 25.5 ± 0.5 | 7.6 ± 0.5 |
| HRas<br>GDPa<sup>*</sup> | 31.7 ± 0.7<br>(1.00 : 0.95) | N/A | 33.3 ± 0.7 | N/A | Not<br>measurable | 33.3 ± 0.7 |
| HRas<br>GDPb<sup>*</sup> | 20.0 ± 0.3<br>(1.00 : 0.94) | N/A | 21.3 ± 0.2 | N/A | Not<br>measurable | 21.3 ± 0.2 |
| HRas<br>GDPC<sup>*</sup> | 40.1 ± 0.3<br>(1.00 : 0.96) | N/A | 41.8 ± 0.4 | N/A | Not<br>measurable | 41.8 ± 0.4 |

## Notes

### Competing Interest Statement

The authors have declared no competing interest.

## References

[1] A.D. Cox, C.J. Der, Ras history: The saga continues, Small GTPases, 1 (2010) 2–27.

[2] K. Wuichet, L. Sogaard-Andersen, Evolution and diversity of the Ras superfamily of small GTPases in prokaryotes, Genome Biol Evol, 7 (2014) 57–70.

[3] I.M. Ahearn, K. Haigis, D. Bar-Sagi, M.R. Philips, Regulating the regulator: post-translational modification of RAS, Nat Rev Mol Cell Biol, 13 (2011) 39–51.

[4] M. Drosten, A. Dhawahir, E.Y. Sum, J. Urosevic, C.G. Lechuga, L.M. Esteban, E. Castellano, C. Guerra, E. Santos, M. Barbacid, Genetic analysis of Ras signalling pathways in cell proliferation, migration and survival, EMBO J, 29 (2010) 1091–1104.

[5] D.K. Simanshu, D.V. Nissley, F. McCormick, RAS Proteins and Their Regulators in Human Disease, Cell, 170 (2017) 17–33.

[6] E.F. Pai, W. Kabsch, U. Krengel, K.C. Holmes, J. John, A. Wittinghofer, Structure of the guanine-nucleotide-binding domain of the Ha-ras oncogene product p21 in the triphosphate conformation, Nature, 341 (1989) 209–214.

[7] M.V. Milburn, L. Tong, A.M. deVos, A. Brunger, Z. Yamaizumi, S. Nishimura, S.H. Kim, Molecular switch for signal transduction: structural differences between active and inactive forms of protooncogenic ras proteins, Science, 247 (1990) 939–945.

[8] A.J. Scheidig, C. Burmester, R.S. Goody, The pre-hydrolysis state of p21(ras) in complex with GTP: new insights into the role of water molecules in the GTP hydrolysis reaction of ras-like proteins, Structure, 7 (1999) 1311–1324.

[9] E.F. Pai, U. Krengel, G.A. Petsko, R.S. Goody, W. Kabsch, A. Wittinghofer, Refined crystal structure of the triphosphate conformation of H-ras p21 at 1.35 A resolution: implications for the mechanism of GTP hydrolysis, EMBO J, 9 (1990) 2351–2359.

[10] A.M. Rojas, G. Fuentes, A. Rausell, A. Valencia, The Ras protein superfamily: evolutionary tree and role of conserved amino acids, J Cell Biol, 196 (2012) 189–201.

[11] R. Gasper, F. Wittinghofer, The Ras switch in structural and historical perspective, Biol Chem, 401 (2019) 143–163.

[12] A.G. Stephen, D. Esposito, R.K. Bagni, F. McCormick, Dragging ras back in the ring, Cancer Cell, 25 (2014) 272–281.

[13] C. Kiel, D. Matallanas, W. Kolch, The Ins and Outs of RAS Effector Complexes, Biomolecules, 11 (2021).

[14] J. John, R. Sohmen, J. Feuerstein, R. Linke, A. Wittinghofer, R.S. Goody, Kinetics of interaction of nucleotides with nucleotide-free H-ras p21, Biochemistry, 29 (1990) 6058–6065.

[15] I.R. Vetter, A. Wittinghofer, The guanine nucleotide-binding switch in three dimensions, Science, 294 (2001) 1299–1304.

[16] B.E. Hall, D. Bar-Sagi, N. Nassar, The structural basis for the transition from Ras-GTP to Ras-GDP, Proc Natl Acad Sci U S A, 99 (2002) 12138–12142.

[17] C. Kotting, K. Gerwert, Time-resolved FTIR studies provide activation free energy, activation enthalpy and activation entropy for GTPase reactions, Chem Phys, 307 (2004) 227–232.

[18] S.E. Neal, J.F. Eccleston, M.R. Webb, Hydrolysis of Gtp by P21nras, the Nras Protooncogene Product, Is Accompanied by a Conformational Change in the Wild-Type Protein - Use of a Single Fluorescent-Probe at the Catalytic Site, P Natl Acad Sci USA, 87 (1990) 3562–3565.

[19] J. Tucker, G. Sczakiel, J. Feuerstein, J. John, R.S. Goody, A. Wittinghofer, Expression of p21 proteins in Escherichia coli and stereochemistry of the nucleotide-binding site, EMBO J, 5 (1986) 1351–1358.

[20] J.G. Tate, S. Bamford, H.C. Jubb, Z. Sondka, D.M. Beare, N. Bindal, H. Boutselakis, C.G. Cole, C. Creatore, E. Dawson, P. Fish, B. Harsha, C. Hathaway, S.C. Jupe, C.Y. Kok, K. Noble, L. Ponting, C.C. Ramshaw, C.E. Rye, H.E. Speedy, R. Stefancsik, S.L. Thompson, S. Wang, S. Ward, P.J. Campbell, S.A. Forbes, COSMIC: the Catalogue Of Somatic Mutations In Cancer, Nucleic Acids Res, 47 (2019) D941–D947.

[21] G. Buhrman, G. Wink, C. Mattos, Transformation efficiency of RasQ61 mutants linked to structural features of the switch regions in the presence of Raf, Structure, 15 (2007) 1618–1629.

[22] C.E. Burd, W. Liu, M.V. Huynh, M.A. Waqas, J.E. Gillahan, K.S. Clark, K. Fu, B.L. Martin, W.R. Jeck, G.P. Souroullas, D.B. Darr, D.C. Zedek, M.J. Miley, B.C. Baguley, S.L. Campbell, N.E. Sharpless, Mutation-specific RAS oncogenicity explains NRAS codon 61 selection in melanoma, Cancer Discov, 4 (2014) 1418–1429.

[23] J.C. Hunter, A. Manandhar, M.A. Carrasco, D. Gurbani, S. Gondi, K.D. Westover, Biochemical and Structural Analysis of Common Cancer-Associated KRAS Mutations, Mol Cancer Res, 13 (2015) 1325–1335.

[24] M.J. Smith, B.G. Neel, M. Ikura, NMR-based functional profiling of RASopathies and oncogenic RAS mutations, Proc Natl Acad Sci U S A, 110 (2013) 4574–4579.

[25] I.A. Prior, P.D. Lewis, C. Mattos, A comprehensive survey of Ras mutations in cancer, Cancer Res, 72 (2012) 2457–2467.

[26] R.C. Gimple, X. Wang, RAS: Striking at the Core of the Oncogenic Circuitry, Front Oncol, 9 (2019) 965.

[27] C.W. Johnson, G. Buhrman, P.Y. Ting, J. Colicelli, C. Mattos, Expression, purification, crystallization and X-ray data collection for RAS and its mutants, Data Brief, 6 (2016) 423–427.

[28] M. Stumber, C. Herrmann, S. Wohlgemuth, H.R. Kalbitzer, W. Jahn, M. Geyer, Synthesis, characterization and application of two nucleoside triphosphate analogues, GTPgammaNH(2) and GTPgammaF, Eur J Biochem, 269 (2002) 3270-3278.

[29] M.E. Pacold, S. Suire, O. Perisic, S. Lara-Gonzalez, C.T. Davis, E.H. Walker, P.T. Hawkins, L. Stephens, J.F. Eccleston, R.L. Williams, Crystal structure and functional analysis of Ras binding to its effector phosphoinositide 3-kinase gamma, Cell, 103 (2000) 931–943.

[30] S.K. Fetics, H. Guterres, B.M. Kearney, G. Buhrman, B. Ma, R. Nussinov, C. Mattos, Allosteric effects of the oncogenic RasQ61L mutant on Raf-RBD, Structure, 23 (2015) 505–516.

[31] R. Qamra, S.R. Hubbard, Structural basis for the interaction of the adaptor protein grb14 with activated ras, PLoS One, 8 (2013) e72473.

[32] B. Stieglitz, C. Bee, D. Schwarz, O. Yildiz, A. Moshnikova, A. Khokhlatchev, C. Herrmann, Novel type of Ras effector interaction established between tumour suppressor NORE1A and Ras switch II, EMBO J, 27 (2008) 1995–2005.

[33] M.G. Rudolph, P. Bayer, A. Abo, J. Kuhlmann, I.R. Vetter, A. Wittinghofer, The Cdc42/Rac interactive binding region motif of the Wiskott Aldrich syndrome protein (WASP) is necessary but not sufficient for tight binding to Cdc42 and structure formation, J Biol Chem, 273 (1998) 18067–18076.

[34] G.H. Swisher, J.P. Hannan, N.J. Cordaro, A.H. Erbse, J.J. Falke, Ras-Guanine Nucleotide Complexes: A UV Spectral Deconvolution Method to Analyze Protein Concentration, Nucleotide Stoichiometry, and Purity, Anal Biochem, (2021) 114066.

[35] T.C. Buckles, B.P. Ziemba, G.R. Masson, R.L. Williams, J.J. Falke, Single-Molecule Study Reveals How Receptor and Ras Synergistically Activate PI3Kalpha and PIP3 Signaling, Biophys J, 113 (2017) 2396–2405.

[36] J. Gureasko, W.J. Galush, S. Boykevisch, H. Sondermann, D. Bar-Sagi, J.T. Groves, J. Kuriyan, Membrane-dependent signal integration by the Ras activator Son of sevenless, Nat Struct Mol Biol, 15 (2008) 452–461.

[37] W.C. Lin, L. Iversen, H.L. Tu, C. Rhodes, S.M. Christensen, J.S. Iwig, S.D. Hansen, W.Y. Huang, J.T. Groves, H-Ras forms dimers on membrane surfaces via a protein-protein interface, Proc Natl Acad Sci U S A, 111 (2014) 2996–3001.

[38] J.G. Martyr, Effects of H-Ras Disease Mutations on Binding to the Ras Binding Domain of PI3KD Measured by Micro-Scale Thermophoresis, Department of Biochemistry, University of Colorado Boulder, https://scholar.colorado.edu/concern/undergraduate_honors_theses/6t053g59n, 2019.

[39] R.C. Killoran, M.J. Smith, Conformational resolution of nucleotide cycling and effector interactions for multiple small GTPases determined in parallel, J Biol Chem, 294 (2019) 9937–9948.

[40] R.M. Bock, N.-S. Ling, S.A. Morell, S.H. Lipton, Ultraviolet absorption spectra of adenosine- 5’-triphosphate and related 5’-ribonucleotides, Archives of Biochemistry and Biophysics, 62 (1956) 253–264.

[41] A. Eberth, M.R. Ahmadian, In vitro GEF and GAP assays, Curr Protoc Cell Biol, Chapter 14 (2009) Unit 14 19.

[42] C. Lenzen, R.H. Cool, A. Wittinghofer, Analysis of intrinsic and CDC25-stimulated guanine nucleotide exchange of p21ras-nucleotide complexes by fluorescence measurements, Methods Enzymol, 255 (1995) 95–109.

[43] I. Rubio, R. Pusch, R. Wetzker, Quantification of absolute Ras-GDP/GTP levels by HPLC separation of Ras-bound [(32)P]-labelled nucleotides, J Biochem Biophys Methods, 58 (2004) 111–117.

[44] P. Bandaru, N.H. Shah, M. Bhattacharyya, J.P. Barton, Y. Kondo, J.C. Cofsky, C.L. Gee, A.K. Chakraborty, T. Kortemme, R. Ranganathan, J. Kuriyan, Deconstruction of the Ras switching cycle through saturation mutagenesis, Elife, 6 (2017) 27810.

[45] F. Paul, H. Zauber, L. von Berg, O. Rocks, O. Daumke, M. Selbach, Quantitative GTPase Affinity Purification Identifies Rho Family Protein Interaction Partners, Mol Cell Proteomics, 16 (2017) 73–85.

[46] M. Jaiswal, B.N. Dubey, K.T. Koessmeier, L. Gremer, M.R. Ahmadian, Biochemical assays to characterize Rho GTPases, Methods Mol Biol, 827 (2012) 37–58.

[47] M. Hanzal-Bayer, L. Renault, P. Roversi, A. Wittinghofer, R.C. Hillig, The complex of Arl2-GTP and PDE delta: from structure to function, EMBO J, 21 (2002) 2095–2106.

[48] Y. Bromberg, E. Shani, G. Joseph, Y. Gorzalczany, O. Sperling, E. Pick, The GDP-bound form of the small G protein Rac1 p21 is a potent activator of the superoxide-forming NADPH oxidase of macrophages, J Biol Chem, 269 (1994) 7055–7058.

[49] S.J.M. Smith, K. Rittinger, Preparation of GTPases for structural and biophysical analysis, in: E. Manser, T. Leung (Eds.) GTPase Protocols: The Ras Superfamily, Humana Press, Totowa, New Jersey, 2002, pp. 13–24.

[50] K.M. Ferguson, T. Higashijima, M.D. Smigel, A.G. Gilman, The influence of bound GDP on the kinetics of guanine nucleotide binding to G proteins, J Biol Chem, 261 (1986) 7393–7399.

[51] O. Weiss, J. Holden, C. Rulka, R.A. Kahn, Nucleotide binding and cofactor activities of purified bovine brain and bacterially expressed ADP-ribosylation factor, J Biol Chem, 264 (1989) 21066–21072.

[52] B. Jadhav, K. Wild, M.R. Pool, I. Sinning, Structure and Switch Cycle of SRbeta as Ancestral Eukaryotic GTPase Associated with Secretory Membranes, Structure, 23 (2015) 1838–1847.

[53] C.H. Gray, J. Konczal, M. Mezna, S. Ismail, J. Bower, M. Drysdale, A fully automated procedure for the parallel, multidimensional purification and nucleotide loading of the human GTPases KRas, Rac1 and RalB, Protein Expr Purif, 132 (2017) 75–84.

[54] A. Contreras-Sanz, T.S. Scott-Ward, H.S. Gill, J.C. Jacoby, R.E. Birch, J. Malone-Lee, K.M. Taylor, C.M. Peppiatt-Wildman, S.S. Wildman, Simultaneous quantification of 12 different nucleotides and nucleosides released from renal epithelium and in human urine samples using ion-pair reversed-phase HPLC, Purinergic Signal, 8 (2012) 741–751.

[55] H.M. Berman, J. Westbrook, Z. Feng, G. Gilliland, T.N. Bhat, H. Weissig, I.N. Shindyalov, P.E. Bourne, The Protein Data Bank, Nucleic Acids Res, 28 (2000) 235–242.

[56] M.A. Gray, S.A. Austin, M.J. Clemens, L. Rodrigues, C.A. Pasternak, Protein synthesis in Semliki Forest virus-infected cells is not controlled by permeability changes, J Gen Virol, 64 (Pt 12) (1983) 2631–2640.

[57] C. Munoz-Maldonado, Y. Zimmer, M. Medova, A Comparative Analysis of Individual RAS Mutations in Cancer Biology, Front Oncol, 9 (2019) 1088.

[58] I. van der Burgt, W. Kupsky, S. Stassou, A. Nadroo, C. Barroso, A. Diem, C.P. Kratz, R. Dvorsky, M.R. Ahmadian, M. Zenker, Myopathy caused by HRAS germline mutations: implications for disturbed myogenic differentiation in the presence of constitutive HRas activation, J Med Genet, 44 (2007) 459–462.

[59] S.A. Moodie, M. Paris, E. Villafranca, P. Kirshmeier, B.M. Willumsen, A. Wolfman, Different structural requirements within the switch II region of the Ras protein for interactions with specific downstream targets, Oncogene, 11 (1995) 447–454.

